# Modelling Osteocyte Network Formation: Healthy and Cancerous Environments

**DOI:** 10.1101/729046

**Authors:** Jake P. Taylor-King, Pascal R. Buenzli, S. Jon Chapman, Conor C. Lynch, David Basanta

**Affiliations:** Institute of Molecular Systems Biology, Department of Biology, ETHZ, CH-8093 Zurich, Switzerland; Mathematical Institute, University of Oxford, Oxford, OX2 6GG, UK; Integrated Mathematical Oncology, H. Lee Moffitt Cancer Center and Research Institute, Tampa, FL, USA; School of Mathematical Sciences, Queensland University of Technology, Brisbane, QLD 4001, Australia; Department of Tumor Biology, H. Lee Moffitt Cancer Center and Research Institute, Tampa, FL, USA

## Abstract

Advanced cancers, such as prostate and breast cancers, commonly metastasize to bone. In the bone matrix, dendritic osteocytes form a spatial network allowing communication between osteocytes and the osteoblasts located on the bone surface. This communication network facilitates coordinated bone remodelling. In the presence of a cancerous microenvironment, the morphology of this network changes. Commonly osteocytes appear to be either overdifferentiated (i.e., there are more dendrites than healthy bone) or underdeveloped (i.e., dendrites do not fully form). In addition to structural changes, histological sections from metastatic breast cancer xenografted mice show that number of osteocytes per unit area is different between healthy bone and cancerous bone. We present a stochastic agent-based model for bone formation incorporating osteoblasts and osteocytes that allows us to probe both network structure and density of osteocytes in bone. Our model both allows for the simulation of our spatial network model and analysis of mean-field equations in the form of integro-partial differential equations. We considered variations of our model to study specific physiological hypotheses related to osteoblast differentiation; for example predicting how changing biological parameters, such as rates of bone secretion, rates of cancer formation and rates of osteoblast differentiation can allow for qualitatively different network morphologies. We then used our model to explore how commonly applied therapies such as bisphosphonates (e.g. zoledronic acid) impact osteocyte network formation.

## Introduction

Advanced cancers commonly metastasize to bone where they often disrupt the normal bone remodelling process [1, 2]. With the onset of various types of bone cancer, it is common for the bone remodelling process to be disrupted. Much previous work has been focused on macroscopic properties of the resulting bone, e.g., the osteoblastic (net bone formation) and osteolytic (net bone reduction) phenotypes. As bone is formed and resorbed cyclically, osteocyte networks can be morphologically malformed. A relatively unexplored area regarding cancerous bone formation is the study of osteoblast-to-osteocyte differentiation whilst concurrently taking into account network structure. Understanding the full nature of bone formation networks is now becoming extremely important, especially as osteocyte network structure is suspected to limit effectiveness of current anti-cancer therapy [3].

Myeloma and (benign) osteoma, osteocytes appear exceptionally spherical with shorter distorted dendrites that are reduced in number [4, 5]. An experimentally contrasting osteocyte network was observed with unregulated excessive cancerous growth in the presence of osteogenic sarcoma [4]. Broadly speaking, osteocytes within a cancerous microenvironment display either over or under developed phenotypes (see Table 1 and figures in Ref. [4]) leading to “more connected” or “less connected” networks.

**Table 1.** Cancer type impact on osteocyte morphology.

| Cancer type | Origin | Bone growth | Osteocyte morphology | Ref. |
| --- | --- | --- | --- | --- |
| Osteogenic Sarcoma. | Mesenchymal cells | Mixture | Canaliculi upregulated, Lacunae empty. | [4] |
| Osteoma (benign). | Unknown | Osteoblastic | Canaliculi down regulated, retarded growth. | [4] |
| Myeloma | Plasma cells / White blood cells | Primarily osteolytic | Osteocyte lacunae spherical, canaliculi reduced in number, shorter, distorted. | [5]<br>[6] |
| Metastasis (group A) | Lung, Breast, Liver | Primarily osteolytic | Unknown | [5] |
| Metastasis (group B) | Prostate | Primarily osteoblastic | Unknown | [7] |

Perturbations in osteocyte-network organization can impact both fluid flow and diffusion of metabolites and thereby affect mechanosensation and signaling [3, 8–10]. The exchange of signalling molecules through the lacuno-canalicular network relate to: skeletal unloading, and fatigue damage [11]. The range of signaling molecules that have been detected is vast and arise in the regulation of bone mineralisation [12], and many other organs^1^. Studies have reported that high-density networks correlate positively with high bone quality [8].

The functional role of these different network topologies is unclear. Experimental works only reveal a snapshot of the communication between the osteocytes within bone, and the osteoblasts on the bone surface. However, one can use mathematical modelling as a tool for investigation. Marotti *et. al*. suggested that osteoblasts are incorporated into the osteocyte network by mature osteocytes extending their dendrites towards the osteoblast layer [16–20]. Experimentally, sclerostin has been stained for and observed within the cancer structures between osteocytes and osteoblasts [21]. These studies highlight the important roleosteocytes play in osteoblast function and differentiation.

Therapeutically, zoledronic acid is frequently used to treat metastatic breast cancer (BCA) where pathological bone is formed with lower densities of osteocytes per unit volume; zoledronate then helps recover the number of osteocytes by inhibiting osteoclasts — although it is not clear if the network structure is restored.

In this paper, we have developed a stochastic agent-based model to investigate how cancer cells regulate osteocyte behaviour and bone formation. We consider osteocyte network formation building on an earlier model of osteocyte generation [22], which did not account for network structure. In said model, osteocytes are located within a growing domain that represents the presence of mineralised bone; osteoblasts are located on the surface of the bone substrate. There are two constitutive processes: (*i*) the bone surface moves with a speed proportional to the surface density of osteoblasts; and (*ii*) osteoblasts differentiate into osteocytes. We add to the model by allowing for extra structure relating to the osteocyte’s canicular network. We allow for osteocytes to extend dendrites towards the osteoblast layer to signal osteoblast differentiation (we assume these processes occur concurrently). Osteoblasts are also allowed to proliferate and move along the bone surface.

With this model, we aim to link osteocyte density and network structure to biological quantities such as: the rate of osteoid secretion; the rate of osteocyte network formation; and the rate of preosteoblast proliferation. In particular, we investigate how stimulatory or inhibitory network dependent signals influence osteoblast-to-osteocyte differentiation and lead to different osteocyte network properties in newly formed bone.

## 1 Materials and Methods

### 1.1 Histological Analysis of Osteocyte Density

Specimens for analysis were derived from mice that were intratibially inoculated with either saline (Control) or breast cancer (BCa, PyMT cell line) as described previously under University of South Florida IACUC approved protocols (R2238 and R1762-CCL)[23, 24]. Mice were harvested for analysis prior to breach of the cortical bone (day 21). Separately, BCa-PyMT inoculated mice were treated with a bisphosphonate (zoledronate) over the course of the study period (1mg/Kg, subcutaneously thrice weekly) [23, 24]. Subsequent to tissue collection and isolation, bones were decalcified, processed and paraffin embedded. Sections (5*μ*m) were generated, rehydrated and then stained with either Gomori’s Trichrome or Hematoxylin & Eosin using standard procedures.

We estimated the area of visible trabecular bone within a pathology image, and subsequently count the number of osteocytes within this region. For full details of this Algorithm, see Appendix A. Unfortunately, the samples were not at a resolution high enough to determine network structure, however it was possible to estimate number of osteocytes per unit area (#osteocytes/mm^2^).These results are presented as box plots in Figure 1.

**Fig 1.**
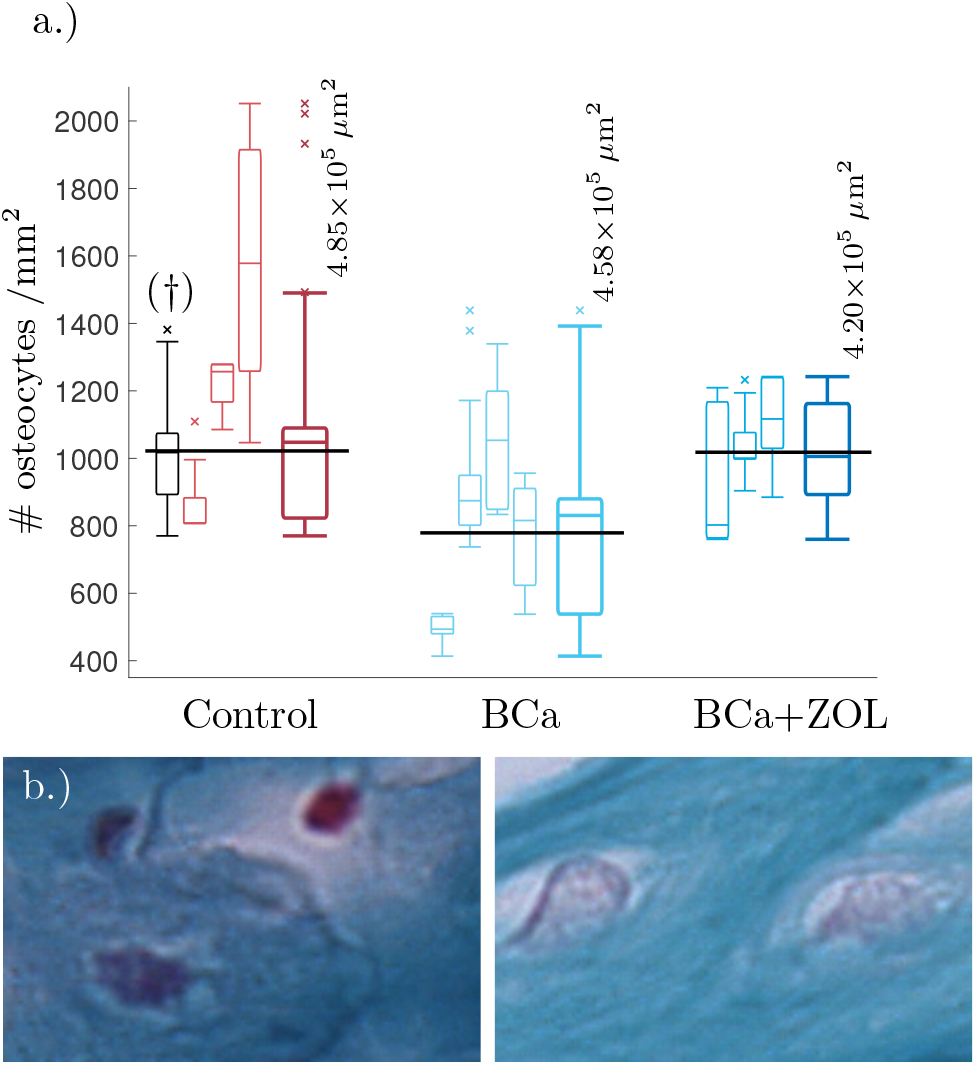
(a) Tukey box plot showing results to preliminary image analysis of osteocyte number density. Means plotted in black (see text for details). The far left histogram (marked †) is representative of a collection of histology slides imaged at a lower resolution. (b) Comparison between osteocyte sizes: (left) control, (right) bone under breast cancer protocol. Image size corresponds to 32 × 21 *μ*m. The smaller box plots are the osteocyte densities for each mouse using 3–8 slides per mouse. Each sample of the osteocyte number density is weighted by the quantity of visible bone area within the sample when determining the data statistics summarised in the plots. The larger box plots show the combined data for mice undergoing the same protocol.

From Figure 1, breast cancer pathological bone have significant lower osteocyte number densities when compared to healthy bone. Macroscopically, breast cancer is often osteolytic and suppresses osteoblast proliferation and maturation. When applied as therapy, the zoledronate treatment allows for recovery of osteocyte number density.

### 1.2 Mathematical Model

#### 1.2.1 Mathematical Description

Our model is a stochastic agent-based model for the bone formation phase only. The agents in our model are osteocytes and osteoblasts occupying nodes within a spatial network. The osteoblast-to-osteocyte differentiation pathway is subdivided into eight phenotypic stages: (*i*) preosteoblast; (*ii*) preosteoblastic osteoblast; (*iii*) osteoblast; (*iv*) osteoblastic osteocyte; (*v*) osteoid-osteocyte (Type II preosteocyte); (*vi*) Type III preosteocyte; (*vii*) young osteocyte; and (*viii*) old osteocyte [25]. Additionally, the secretion of bone occurs as two steps: first osteoid is deposited as a collageneous scaffold, and then mineralization occurs to confer strength. Stages (*iv*)–(*vi*) are cells after the deposition front but before the mineralization front, surrounded by a non-mineralized osteoid matrix (i.e., there is scaffold around them). Stages (*vii*)–(*viii*) are cells whose volume has depleted (endoplasmic reticulum and Golgi apparatus reduction) and are in mineralised bone. The diagram in Fig. 2 shows the bone-formation step. Here we are only interested in the structure of a mature osteocyte network [stages (*vi*)–(*viii*)], so we avoid modelling the full biology intricacy (e.g., cell sub-classifications, proteins etc) for simplicity. We model mobile preosteoblasts (disconnected from the osteocyte network) in front of the bone deposition front that proliferate; stationary mature osteoblasts that secrete osteoid and are connected to the osteocyte network; and osteocytes that form dendrites with mature osteoblasts.

**Fig 2.**
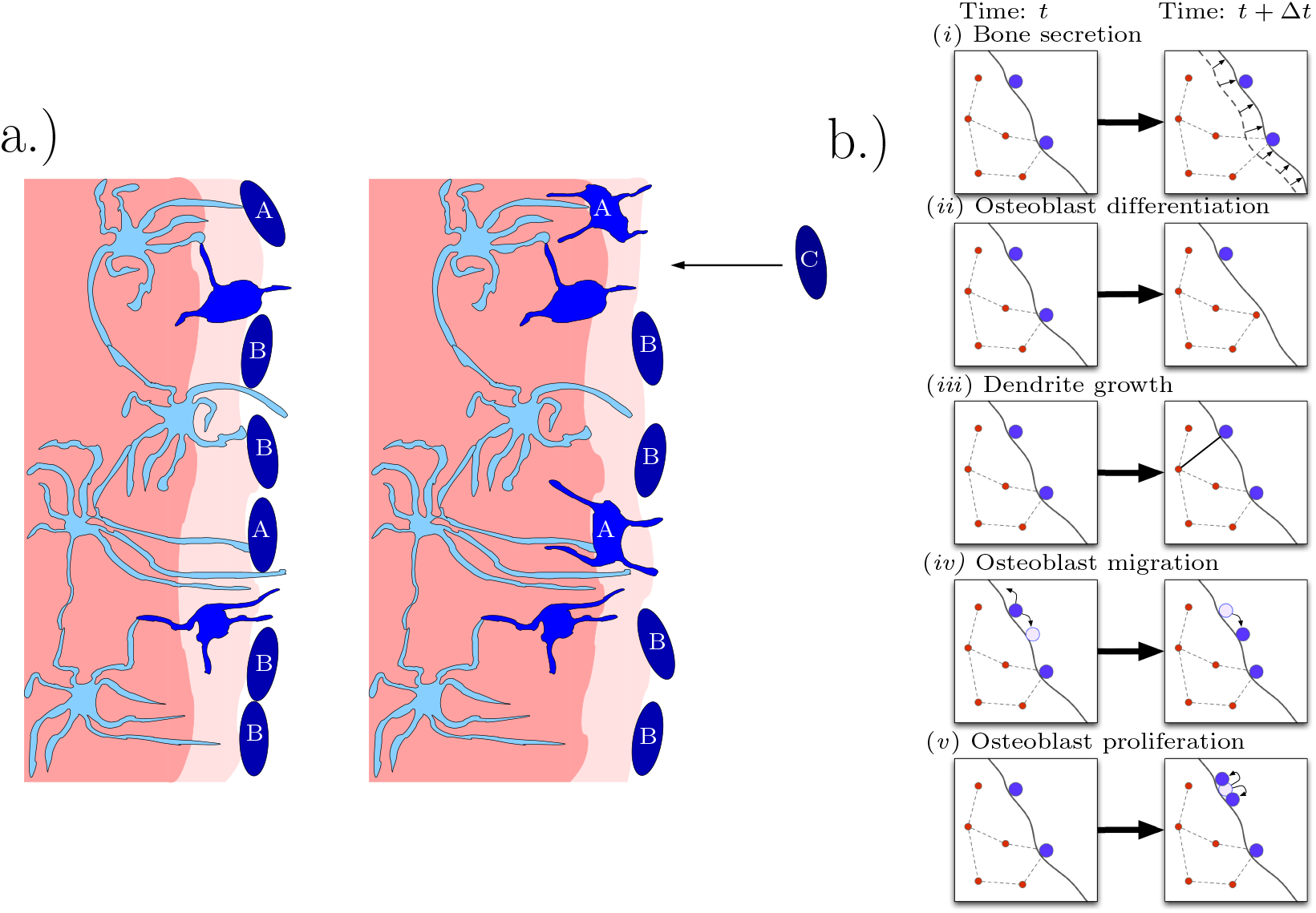
Diagrammatic illustration of bone formation. (a.) Biological process. Lighter shades of blue indicate more differentiated cells. The lighter shade of pink indicates the deposition front, and the darker shade of pink indicates the mineralization front. The left panel occurs earlier than the right panel. Dendritic osteocytes (light blue) have dendrites that extend towards the osteoblast layer (dark blue). The osteoblasts secrete bone matrix. Osteoblast cells marked with “A” are signaled by the osteocyte network to differentiate into osteocytes. Osteoblast cells marked with “B” do not differentiate and stay on the outer bone surface. Osteoblast cells marked with “C” arrive at the bone front after differentiating from precursor osteoblasts (preosteoblasts). (This figure is adapted from Ref. [25].) (b.) Model representation of biological process. In a small time step, the following events can occur: (*i*) bone secretion; (*ii*) osteoblast differentiation; (*iii*) dendrite growth; (*iv*) osteoblast migration; and (*v*) osteoblast proliferation.

In our model, the spatial network has pairwise edges between cells, representing the ability for two cells to communicate; physically this communication is mediated through dendrites.

The model consists of the following processes: (*i*) bone secretion; (*ii*) osteoblast differentiation; (*iii*) dendrite growth; (*iv*) preosteoblast migration; and (*v*) preosteoblast proliferation (see Fig. 2). Agents of the same type (osteocytes or osteoblasts) follow the same rules, although each agent may have different properties, e.g., position, number of edges, in addition to type.

Bone secretion is carried out by osteoblasts secreting bone in a small region around themselves, orthogonal to the bone surface (in the normal direction). Osteoblasts can also become buried and differentiate into osteocytes. The rate of osteoblast differentiation may depend on if the osteoblast in question is in communication with the osteocyte network in bone.

For dendrite growth, osteocytes and osteoblasts create a communication channel (i.e., become connected in the network) at a rate that is a function of the distance between the two cells. Our model is consistent with the suggestions in Refs. [26] that dendrites grow away from osteocytes within bone and towards the osteoblast layer on the bone surface.

Movement and cell division are included for pre-osteoblasts that are disconnected from the osteocyte network. Pre-osteoblasts move along the bone bone surface by means of a diffusive process; they are also able to proliferate and divide into create two daughter cells.

Our simulations are carried out in two dimensions, but they represent a three dimensional slice, in which the third dimension *L_z_* is the typical distance between osteocyte centres (*L_z_* ≈ 40*μ*m), projected onto two dimensions (see Appendix Fig. 7). The *x*-direction is the main direction of bone growth, and the volume occupied by the bone increases in time. We impose periodicity in the *y*-direction (orthogonal to bone growth) to avoid boundary effects.

In all the simulations in this paper, we consider bone formation after a cement line has been deposited. A cement line is a 1-5 *μ*m region of hypermineralised (and collagen deficient) bone [27] is deposited after bone resorption to prepare surfaces for new bone formation. When this cement line is deposited, osteocytes from deep within the bone are not necessarily in communication with osteoblasts on the other side of the cement line [28].

Accordingly, we use the initial condition that there are no old osteocytes to communicate with (no initial network structure at the onset of bone formation) and there is an initial surface density of 6 × 10^3^ mm^−2^ pre-osteoblasts that do not have any network structure. We also specify that once an osteoblast is buried, a new osteoblast takes its place (so the total number of osteoblasts at any time is constant). This new osteoblast has no network structure. Therefore, one can interpret this configuration of the model as the scenario in which pre-osteoblasts are in abundance at the bone interface. New osteoblasts then move into the cell gaps on the bone surface as space becomes available (fig. 3).

**Fig 3.**
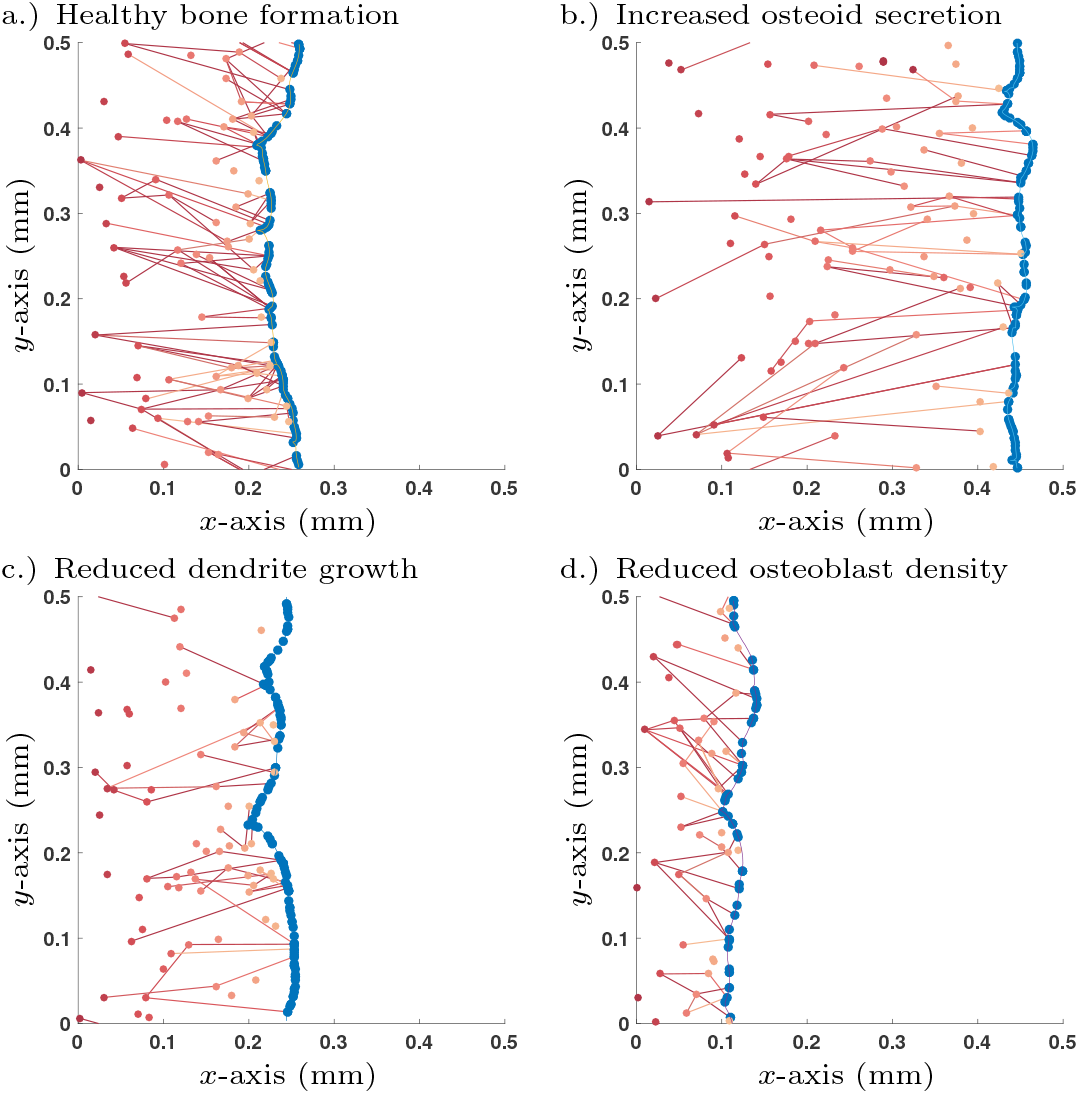
Single simulation runs of bone growth with different parameter choices: a.) Healthy parameter set (as shown in Table 2); b.) Increased osteoid secretion (*η* → 2*η*); c.) Reduced dendrite growth (*α* → *α*/2); d.) Reduced osteoblast surface density 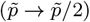. Osteoblast are colored in blue and osteocytes (and their network connections) are in red; darker shades of red denote osteocytes that were buried earlier in time. All simulations are shown after 365 days.

#### 1.2.2 Simulation and Analysis

Individual simulation runs of our stochastic agent-based model were performed with a fixed time step Monte Carlo algorithm (see Appendix B). However, a single realisation is unrepresentative of the stochastic process, so many simulations must be carried out to work out statistics of the process along with parameterisation. This can be computationally demanding, especially as the system grows in size.

As an alternative to large numbers of simulations, we can utilise mean field equations. We have shown previously [29] that in the limit as the number of nodes in the network increases, one is able to derive a mean-field partial integro-differential equation for the expected number of nodes at a particular position in the domain connected to a fixed number of cells. By solving the mean-field equations, one can also calculate the degree distribution of the nodes of the network. The validity of the mean-field assumption is discussed in [29].

Simulations of the stochastic mathematical model suggest that the system is approximately homogeneous in *y*, the direction orthogonal to bone growth (see Fig. 3). Under the assumption of homogeneity in *y*, the mean-field equations depend only on *x*, the direction of bone growth. The solutions of these equations provide spatial profiles of the densities of osteocytes and their network structure perpendicularly to the bone surface (see Appendix C).

Our mean-field equations take the form of two coupled hierarchies of integro-partial differential equations [see equations (23)–(24)]. These equations can be solved to indicate how many particles one could expect to find within a small region of space are connected to *k* other cells (referred to as degree *k*) at time *t*. We denote by *v_k_* the surface density of osteoblasts (number per unit area, [*v_k_*] = mm^−2^) connected to *k* osteocytes on the bone surface; and *w_k_* the number density osteocytes (number per unit volume, [*w_k_*] = mm^−3^) connected to *k* osteocytes or osteoblasts with position *x* at time *t* within the bone. Therefore, we write the total density of all osteoblasts (regardless of degree) as *p* = Σ_*k*_ *v_k_* and total density of osteocytes (regardless of degree) as *q* = Σ_*k*_ *w_k_*. The average number of osteocytes connected to an osteoblast is the average degree of osteoblasts, i.e.,

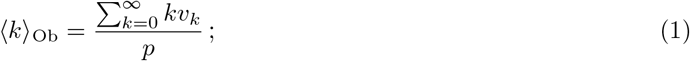

and the average number of cells (osteoblasts and osteocytes) in communication with an osteocyte is the average degree of osteocytes, i.e.,

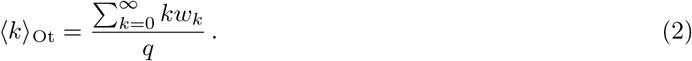

The full equations for *v_k_* and *w_k_* are derived in Appendix C and solved in Appendix D. In the present work, we only consider an uninterrupted bone formation process so that the density of osteocytes *q* at point *x* corresponds to that generated by the terminal differentiation of osteoblasts when the bone deposition front was at *x*.

The mean-field equations admit a travelling wave solution, corresponding to steady bone growth if the osteoblast proliferation rate is such that the surface density of osteoblasts is constant. Such a solution is useful both to explain the model and to investigate the qualitative effects of perturbations to parameters. In the following, we denote the travelling wave solution with a tilde.

## 2 Results

### 2.1 Selection of Differentiation Mechanism

We wish to investigate the effects of different model choices for the rate of osteoblast-to-osteocyte differentiation, *D_k_*.

For a given surface density of osteoblasts, the volumetric density of inclusions embedded in a tissue during bone formation is determined by two dynamic processes: the rate of osteoblast terminal differentiation and the tissue growth rate [22]. If the rate of osteoblast terminal differentiation is identical for all osteoblasts at a given location, i.e., *D_k_* is independent of 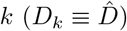, the density of osteocytes generated is given by [22]

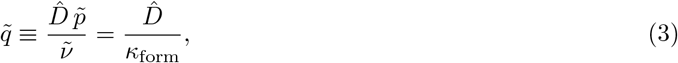

where 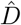 is the terminal differentiation rate, i.e., the probability per unit time for a single osteoblast to become embedded as an osteocyte, 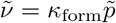 is the normal velocity of the bone interface, and *κ*_form_ is the rate of bone deposition per osteoblast. If the rate of osteoblast terminal differentiation depends additionally on the number of connections *k* they have with osteocytes, the density of osteocytes generated sums up the contributions of all *k*-degree osteoblasts

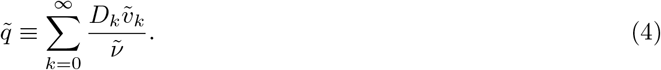

In contrast to equation (3), the density of osteocytes given by equation (4) depends explicitly on the population of osteoblasts, through the proportions 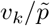 of *k*-degree osteoblasts. Notice that this expression determines the density of osteocytes created at the moving bone deposition front. It does not account for processes that may subsequently affect osteocyte density such as osteocyte apoptosis, bone resorption and remodelling, which may remove and replace osteocytes subsequently [3].

To explore different differentiation mechanisms we alter both: the rate of dendritic growth, *α*, and the mechanism behind the rate of osteoblast differentiation, *D_k_*.

#### 2.1.1 Assuming no network influence: Degree-independent-rate model

To account for the potential of network independent differentiation we assume that the number of osteocytes each osteoblast is in communication with does not impact the rate of osteoblast differentiation; we write

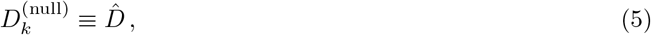

for all values of *k* = 0, 1,…, ∞.

After parameterising the system using experimental measurements found from literature (see Table 2), with this choice of osteoblast differentiation there remains two undefined (free) parameters in the model: the rate of dendrite growth 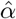, and the rate of osteoblast differentiation 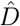; and we calibrate these parameters based on two further experimental observations: the mean degree of an osteoblast connectivity 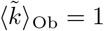 (see Appendix E.6); and the number density of osteocytes 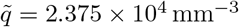. By carrying out mathematical analysis in the travelling wave regime, we obtain formulas linking these quantities together (see Appendix D.2). We determine parameters as 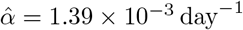 and 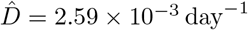.

**Table 2.**
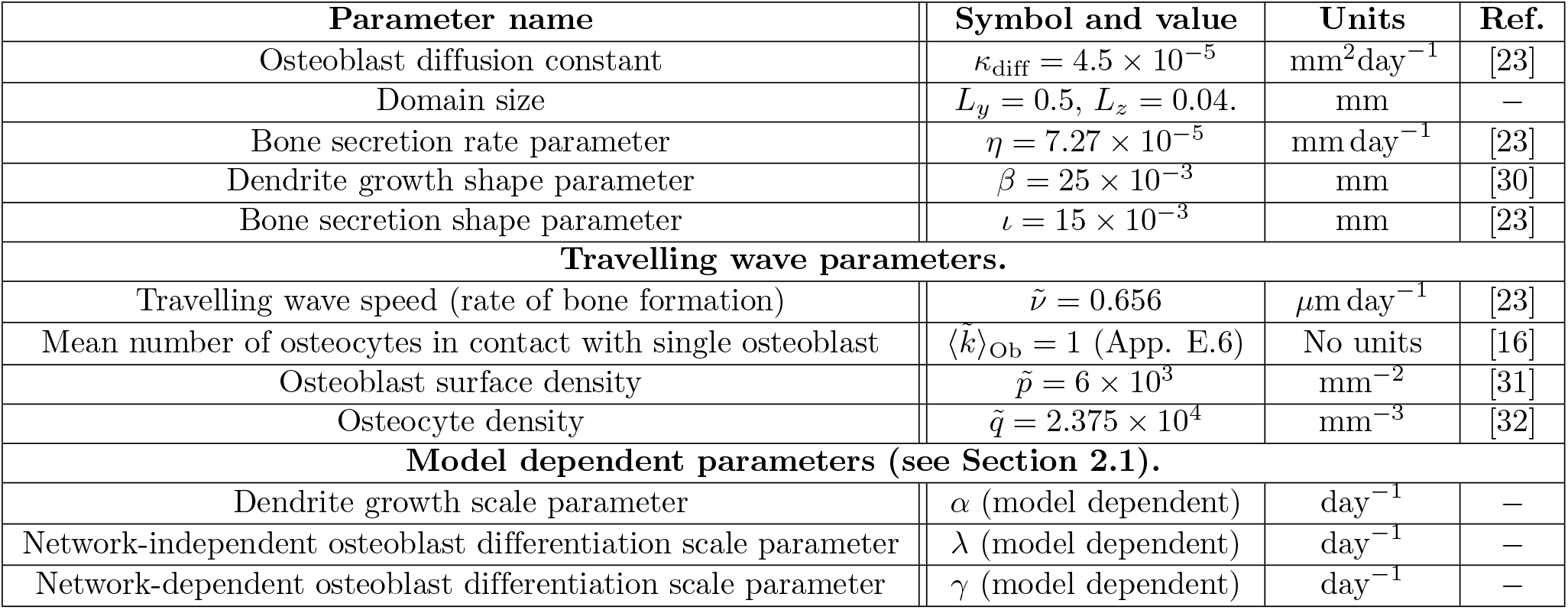
Model parameters, see Appendix E for more details.

If one changes the rate of osteoblast-to-osteocyte differentiation, 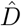, we observe a linear relationship with the number density of osteocytes buried, 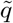 [see equation (3)]. Changing the rate of formation, 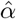, leads to a linear relationship with the mean osteoblast degree, 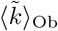, and the mean osteocyte degree, 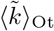. These effects are decoupled — changing the rate of osteoblast-to-osteocyte differentiation, 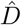, has no effect on the network structure, and changing the rate of dendrite formation, 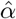, has no effect on the osteocyte density, 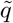. This will not be the case when osteoblast terminal differentiation is coordinated by the osteocyte network.

The assumption that osteoblast differentiation is independent of the osteocyte network is unlikely. First, as reviewed in [33], osteoblasts can be signalled by osteocytes to adhere to the mineral matrix and grow dendrites (subsequently differentiating) via the insulin-like growth factor 1 (IGF-1). Second, sclerostin secreted by osteocytes has been shown to act briefly as an inhibitory signal to prevent excessive osteoblast differentiation and allowing for coordinated osteocyte network formation [21]. Note that in Ref. [21], sclerostin was stained for and observable within the osteocyte’s dendritic protrusions. These experiments, along with the three-dimensional scans taken by Kamioka *et. al*. [16], strongly suggest that osteocytes signal osteoblast differentiation through the extension of dendrites towards the osteoblast layer.

#### 2.1.2 Modelling network effects

We now consider a range of models in which the pre-existing osteocyte network can either have an stimultory or inhibitory effect on osteoblast differentiation. We consider differentiation rates of the form

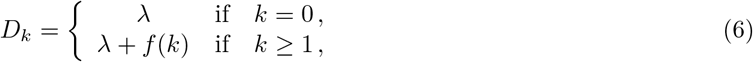

where λ is the network-independent rate of osteoblast differentiation and *f* = *f*(*k*) is the contribution to osteoblast differentiation for an osteoblast connected to *k* osteocytes. When *f* > 0 the network has an excitatory effect on osteoblast differentiation, and when *f* < 0 the network has an inhibitory effect on osteoblast differentiation. To prevent negative differentiation rates, we require that

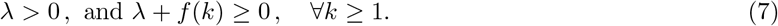

In the main text (see Section 2.1.3 below), we consider only the case where *f* is a constant, *f* = *γ*. Other choices of *f* are considered in Appendix F, i.e., i.e., cumulative activation (*f* ∝ *k*) and diminishing activation (*f* ∝ 1/*k*). It should be noted that choices of *f* that have non-monotonic behaviour (i.e., a local maximum/minimum exists) can lead to non-monotonic profiles in *q*.

#### 2.1.3 Proposed mechanism: Switch-like influence

Switch-like mechanisms are frequently found in biology [34]; at a cellular level this includes initiating mechanisms for proliferation and differentiation [35]. For a switch-like osteoblast differentiation, we take

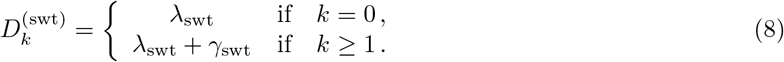

If the osteoblast on the bone surface is not in communication with any other osteocytes (*k* = 0) then it has an network-independent differentiation rate λ_swt_. If osteoblasts are in contact with one or more osteocytes, then there is an induced differentiation rate λ_swt_ + γ_swt_ where γ_swt_ is the added contribution from the dendritic network.

Our technique for parameter identification with this family of terminal differentiation rates is based on the mean-field equations. We choose a network independent component of osteoblast differentiation such that 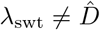, leaving 2 free parameters *α*_swt_ and *γ*_swt_. We then determine the free parameters by imposing that 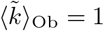 and 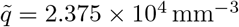 in the travelling wave regime. By making these assertions, if 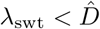, the network contribution will always have an stimulatory effect on osteoblast differentiation; and if 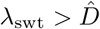, then the network contribution will always inhibit osteoblast differentiation.

To investigate the role of intrinsic to extrinsic osteoblast differentiation rates, we assume that the network-independent differentiation rate λ_swt_ contributes half the total rate of osteoblast-to-osteocyte differentiation rate of the null model (so 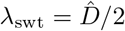). We then determine that *α*_swt_ = 2.08 × 10^−3^ day^−1^ and γ_swt_ = 2.59 × 10^−3^ day^−1^. Another option would be for an inhibitory contribution, in which case we can set 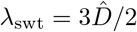, and then *α*_swt_ = 6.96 × 10^−3^ day^−1^ and γ_swt_ = −2.60 × 10^−3^ day^−1^.

#### 2.1.4 Differentiation Mechanism Comparison

In Figure 4, we plot the osteocyte density profile, the mean osteoblast degree over time, and the mean osteocyte degree over time using equations (23)–(25) for 3 proposed choices of *D_k_*: the null model, stimulatory network contributions, and inhibitory network contributions.

**Fig 4.**
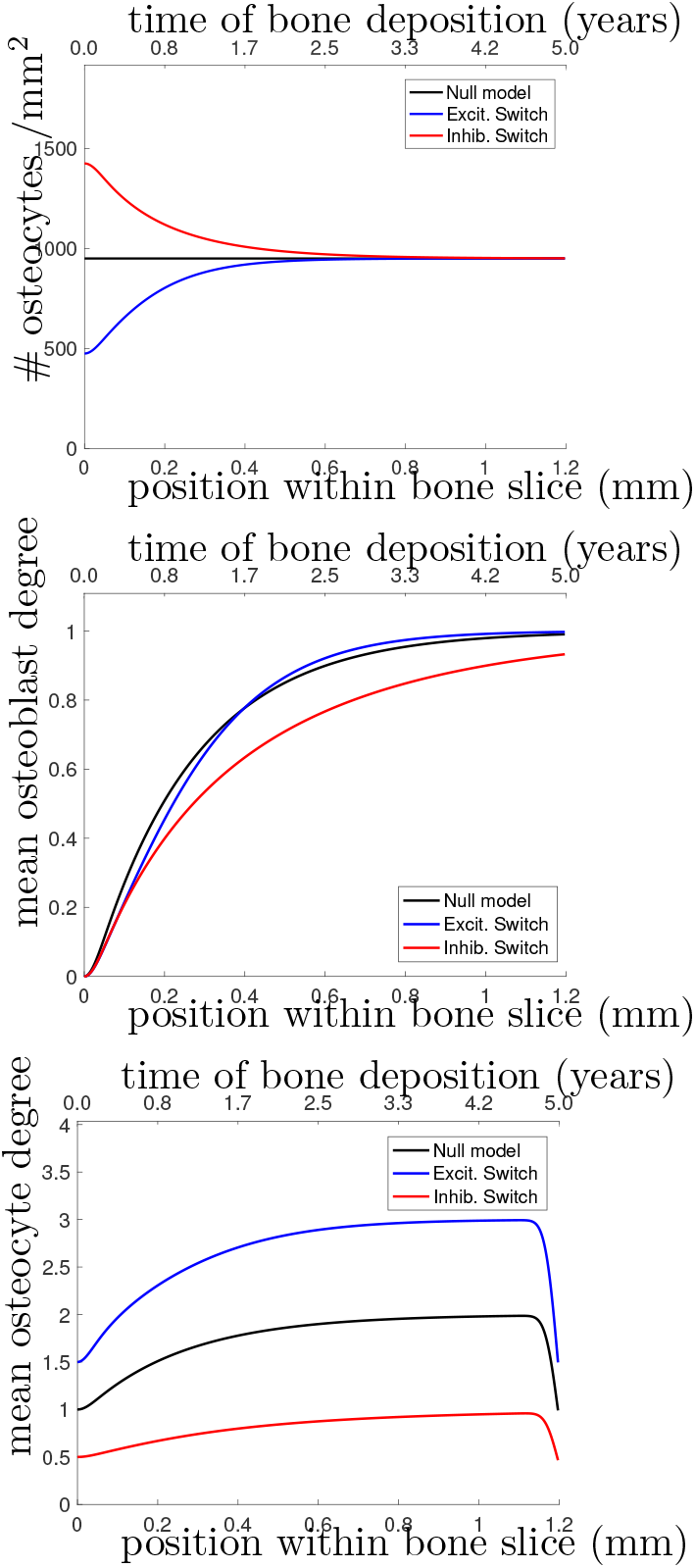
(Top) Osteocyte density profile 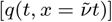, (Middle) mean osteoblast degree over time 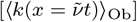, and (Bottom) mean osteocyte degree over time 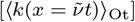 when solving equations (19)–(20). The black line shows the null model, the blue line shows the stimulatory switch model, and the red line shows the inhibitory switch model.

Figure 4 shows that in all 3 models, it takes approximately 1 year (~ 0.7mm) to get to the steady-state desired osteocyte density. We see similar results for the mean osteoblast degree, except that inhibitory contributions to osteoblast differentiation require more time for the osteoblasts to reach the steady-state degree distribution. The main means for differentiation between the three models is the resulting mean osteocyte degree profile. The osteocyte density profile also differs at the onset, but slowly relaxes to the steady state profile. Keeping 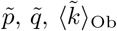 fixed, stimulatory network contributions to osteoblast differentiation leads to more connected osteocyte networks with initially low numbers of osteocytes (higher final value of 〈*k*〉_Ot_), and inhibitory network contributions lead to less connected networks with initially high numbers of osteocytes (lower final value of 〈*k*〉_Ot_).

Capturing network structure and choosing a model that best reflects reality is difficult. For instance, the mean osteoblast degree profile takes a long period of time before settling to the steady-state mean osteoblast degree of 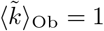. One assumption we have made throughout our model selection process was that the osteoblast surface density was approximately constant. It may be the case that this surface density changes over time [36, 37]. Were it the case that the surface density of osteoblasts was initially higher density before decreasing it would speed up the timescales until a steady-state travelling wave profile was reached.

### 2.2 Parameter Analysis

We explored a wide range of model parameters to further understand their impact on osteocyte network morphology. For the switch-like model in Section 2.1.3, we use the travelling wave analytic expressions described in appendix D.2 to investigate how dependent variables 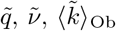, and 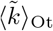 change due to a single perturbation in one of the independent variables *η*, 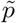, *κ*_diff_, λ, *γ* and *α*, i.e., a sensitivity analysis, see Table 3. In addition to the dependent variables, we also include the number of dendrites per unit area 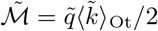 which is a quantity that could be determined experimentally. Depending on whether osteocytes activate or inhibit the rate of osteoblast differentiation, we obtain different results for parameters λ, *γ* and *α*. For visualisation purposes, we show a few stochastic simulation runs for a few specific examples in Figure 3; these examples use the switch-like model in described in section 2.1.3 with the inhibitory parameter configuration (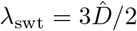, *α*_swt_ = 6.96 × 10^−3^day^−1^, *γ*_swt_ = −2.60 × 10^−3^day^−1^).

**Table 3.** Prediction summary. In the leftmost column, we list parameters that are to be increased. We use the notation that (↑) denotes an increase, (↓) denotes a decrease and (−) signifies no change. Decreasing the parameters in the leftmost column will reverse the directions of the arrows.

| Increased parameter (↑) | Symbol | $\tilde{q}$ | $\tilde{\nu}$ | $\langle \tilde{k} \rangle_{\text{Ob}}$ | $\langle \tilde{k} \rangle_{\text{Ot}}$ | $\tilde{\mathcal{M}}$ |
| --- | --- | --- | --- | --- | --- | --- |
| Rate of bone secretion | $\eta$ | ↓ | ↑ | ↓ | ↓ | ↓ |
| Surface osteoblast density | $\tilde{p}$ | $-$ | ↑ | $-$ | $-$ | $-$ |
| Osteoblast migration speed | $\kappa_{\text{diff}}$ | $-$ | $-$ | $-$ | $-$ | $-$ |
| **Excitatory switch model configuration.** |  |  |  |  |  |  |
| Network-independent rate of osteoblast differentiation | $\lambda$ | ↑ | $-$ | ↑ | $-$ | ↑ |
| Network-dependent rate of osteoblast differentiation | $\gamma$ | ↑ | $-$ | ↓ | $-$ | ↑ |
| Rate of dendrite growth | $\alpha$ | ↑ | $-$ | ↑ | ↑ | ↑ |
| **Inhibitory switch model configuration.** |  |  |  |  |  |  |
| Network-independent rate of osteoblast differentiation | $\lambda$ | ↑ | $-$ | ↓ | $-$ | ↑ |
| Network-dependent rate of osteoblast differentiation | $\gamma$ | ↑ | $-$ | ↓ | $-$ | ↑ |
| Rate of dendrite growth | $\alpha$ | ↓ | $-$ | ↑ | ↑ | ↑ |

Osteocyte density is mostly determined by the secretory rate and terminal differentiation rate, with some weak osteoblast density dependence via the frequency distribution of osteoblast degrees via equation (4). Table 3 confirms this as 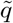 is dependent on the secretory rate (*η*) or the terminal differentiation rate (λ, *γ*, *α*), but not osteoblast density 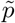. Contrasting these results to those given in Figure 4, Table 3 shows intra-model variability by perturbing parameters, but Figure 4 shows inter-model differences by changing the mechanism for osteoblast differentiation.

Whilst many parameters change the density of osteocytes, only the rate of dendrite growth *α* and the rate of bone formation *η* can change the mean osteocyte degree in the steady state regime. A counter intuitive result also specifies that altering the rate(s) of osteoblast differentiation (parameters λ_swt_ and *γ*_swt_) changes the osteocyte density (variable 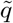) and the osteoblast degree (variable 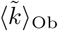), but not the degree of these newly formed osteocytes (variable 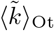). This is possible because osteoblasts are constantly being replaced in the steady state regime; therefore parameters λ_swt_ and *γ*_swt_ are changing the osteocyte density by modifying the mean time osteoblasts secrete bone before differentiation — without changing the mean osteocyte degree.

As we mentioned in 2.1.1, inhibitory network contributions to osteoblast differentiation is more likely in the presence of sclerostin, but stimulatory network contributions may also be possible via IGF-1. Determining which of these two mechanisms drives osteoblast differentiation will require further experimental work.

## 3 Discussion

### 3.1 General implications for cancerous bone growth

A number of previous mathematical models have examined osteocyte density, but none of them have explored network structure. References [38–40] give ordinary differential equation (non-spatial) models for cell populations; these include osteoblast, osteoclast, and osteocyte populations. Existing models of healthy bone remodeling (homeostasis) include spatiotemporal models [41–43], but do not explicitly include osteocyte generation; these models have been adapted for the cancerous regime in Ref. [44]. For TGF*β* targeted therapy, modelling approaches have been used to optimise the treatment window of application [45].

Mechanical focused methods capturing stresses and strains on the bone have also been explored [46, 47]. A general continuum modelling approach was also proposed in [22].

Our model-derived results show that osteocytes are either over differentiated (excessive dendrite growth) or underdeveloped (diminished dendrite growth). Additionally, we show that the osteocyte number density tends to decrease.

With experimental data that gives information on network structure (e.g., transmission electron microscopy, India ink histology stains), one should be able to approximately measure at least 2 of 3 quantities: the number of osteocytes present (quantitative estimate); whether the osteocytes are over-differentiated or underdeveloped (qualitative estimate); and finally the density of dendrites (quantitative estimate). Therefore, we should be able to compare a pathological bone slide to a (healthy) control slide and determine differences between: the osteocyte number density (*q*); the mean osteocyte degree 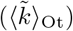; and the density of dendrites 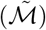.

Furthermore, our model also makes testable predictions in regards to:

a. If all 3 quantities (osteocyte number density, mean osteocyte degree, and dendrite area) have increased (resp. decreased), this corresponds to either: osteoblasts on the bone surface producing too little (resp. too much) osteoid when compared healthy bone; or that the rate of dendrite growth has increased (resp. decreased) in the excitatory switch model configuration.
b. If the osteocyte number density has increased (resp. decreased) with an opposing decrease (resp. increase) in 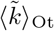 or 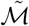, then this must correspond to the rate of dendrite growth changing but in the inhibitory switch model configuration.
c. If the osteocyte density increases (or decreases), but the mean osteocyte conectivity remains constant, our model suggests this relates to a change the rate of osteoblast differentiation.

### 3.2 Implications for Zoledronate Treatment

Breast cancer is known to be osteolytic by promoting osteoclast mediated bone destruction. When applied as a therapy for breast cancer, the zoledronate treatment slows down bone resroption (along with other effects [48]). This treatment is also associated with a recovery of osteocyte number density, see Figure 1.

Changing the proliferation ability of osteoblasts in the model changes the osteoblast surface density. This changes the quantity of bone produced per unit time, but not the network structure or steady state osteocyte density. In our model, changing the process of osteoblast/osteocyte maturation corresponds to changing either: the mechanism behind osteoblast differentiation (parameters λ, *γ*), or changing the rate of dendrite growth (parameter *α*).

In breast cancer (BCa), osteocytes have fewer dendritic connections to other osteocytes and osteocyte density is lower. Therefore the mean osteocyte degree of connectivity is reduced. To achieve a simultaneous decrease in osteocyte density and mean osteocyte degree in the model, one would either have to: decrease the rate of dendrite growth in an stimulatory switch model configuration; or increase the rate of bone secretion per osteoblast.

Given that the zoledronate treatment restores osteocyte density, we propose that future studies investigate how zoledronate acts on the osteoblast-to-osteocyte differentiation pathway. If bone treated with zoledronate has a healthy osteocyte network present, then one can assume zoledronate also targets (and restores) the rate of dendrite growth. However, if bone treated with zoledronate has a different network morphology, one can conclude the differentiation mechanism is targeted, i.e., the network-independent or network-dependent rate of osteoblast differentiation has changed.

## 4 Discussion

We have presented a model for the formation of an osteocyte network and identified parameters for healthy bone formation. By perturbing parameters, one can investigate irregular bone formation and the resulting osteocyte morphology changes. One can also then predict the driving differentiation markers that osteoblasts exhibit that would lead to these morphological changes. In the context of zoledronate therapy for breast cancer, we have used our model to propose how this commonly used clinical treatment impacts bone formation. For future experiments, we have suggested how measurable quantities link to underlying mechanisms.

The model proposed has some limitations, one could suggest many improvements to the model to improve our idealisation of the osteocyte network; we discuss some of these below. Additionally, one might also want to consider the inclusion of chemical species representing proteins of interest, and the inclusion of osteoclasts to incorporate bone resorption. However, our model acts as a first step towards mathematically modelling osteocyte network formation, and avoids making overly specific assumptions on underlying mechanisms.

We now comment on how various aspects of our model compare to biological reality.

### 4.0.1 Triggering osteoblast differentiation

During a bone remodelling event, the total number of osteoblasts generated is far larger than the total number of new osteocytes generated [49]. Pazzaglia *et. al*. [50] have estimated that only 1 in 67 osteoblasts become embedded in bone matrix as osteocytes over the depth of a single osteocyte. In our model using the steady state regime with the parameters from Table 2, over a length-scale of 5 *μ*m approximately 1 in 50 osteoblasts achieve terminal differentiation. The exact mechanisms behind this process are still poorly understood; they may involve physical processes such as burial by neighbouring osteoblasts, or self-burial [25]. It has also been suggested that subpopulations of osteoblasts are predestinated to become osteocytes [51], and that this selection may be determined by the number of connections with osteocytes [16].

One aspect ignored in our model is mechanotransduction. It is known that mechanical loads and fluid flow sheer stress can lead to greater dendrite growth [52, 53]. After now posing our model, a pertinent question then remains as to the relative effect sizes between mechanical stimuli and microenvironmental signalling.

### 4.0.2 Osteocyte degree distribution

In the steady-state travelling wave regime, one can show that the node degrees of the osteocyte network are geometrically distributed when using either the null model (see Section 2.1.1), or the switch-like proposed mechanism (see Section 2.1.3). This effect comes from the difference equation structure shown in (23)–(24) in Appendix C.1.

In Ref. [8] a three-dimensional osteocyte network was studied, the topology of this network includes dendrites that do not connect to a second osteocyte, and edges that link between multiple osteocytes. Additionally, the nodes of their network included both osteocytes and the branching points of dendrites.

Thus the functional communication network we studied is different from the lacuno-canalicular pore network and does not account for all types of communication redundancies that may exist. To incorporate all possible redundancies would require the inclusion of edges that do not connect to other nodes, and edges that exist between multiple nodes. We have some redundancy in our communication network in the form of multiedges (multiple edges between two nodes). However in the limit of large networks, the probability of a multiedge occurring in our model approaches zero. Even in the case of finite networks, this is very unlikely to occur in the model as 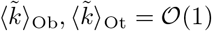. Regardless of these model technicalities, it should be noted that the degree distribution of the three-dimensional scanned network in Ref. [8] was also shown to be geometrically distributed as derived from our model.

### 4.0.3 Orientation of dendrites

It is clear from Figure 3 that we observe orientation of edges between older osteocytes and younger osteocytes or osteoblasts. This leads to an interesting question is as to whether one can configure our model to modify this orientation.

A possible future direction in our work may be to modify our model to explore the functional difference between lateral connections, and connections perpendicular to the bone surface. As osteocytes have coordinated deposition, one may be able to explore whether coordinated terminal differentiation can occur as a function of lateral connections, burying a group of osteoblasts simultaneously.

## Acknowledgments

JPT-K acknowledges funding from the EPSRC (EP/G037280/1). PRB gratefully acknowledges the Australian Research Council for a Discovery Early Career Research Fellowship (project number DE130101191). DB and CCL acknowledges funding from the National Cancer Institute (U01 CA202958-01). We thank Andrew Dhawan and Mason A. Porter for helpful discussions.

## A Image processing technique for Osteocyte density calculation

Using the inbuilt functions of the MATLAB image processing toolbox, our technique for calculating approximate osteocyte densities consists of three stages:

1. Colour-based segmentation to isolate the mineralised section of the bone.
2. Entropy-based segmentation to isolate and locate osteocytes centers.
3. Estimation of osteocyte density as

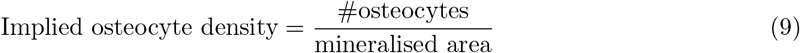

Colour-based segmentation is carried out by treating the RGB signal as a vector and clustering these vectors using *k*-means clustering, the value of *k* chosen depending on an *ad-hoc* basis depending on the specific image. Entropy-based segmentation and thresholding was used to detect approximate locations of osteocytes. Manual validation was also carried out to confirm/correct the locations of the osteocytes detected. In Figure 5, we see the output of the algorithm, the stroma appears red, the mineralised region of the bone appears blue. The region between measured is outlined in green and the osteocyte’s center is marked with a red circle.

**Fig 5.**
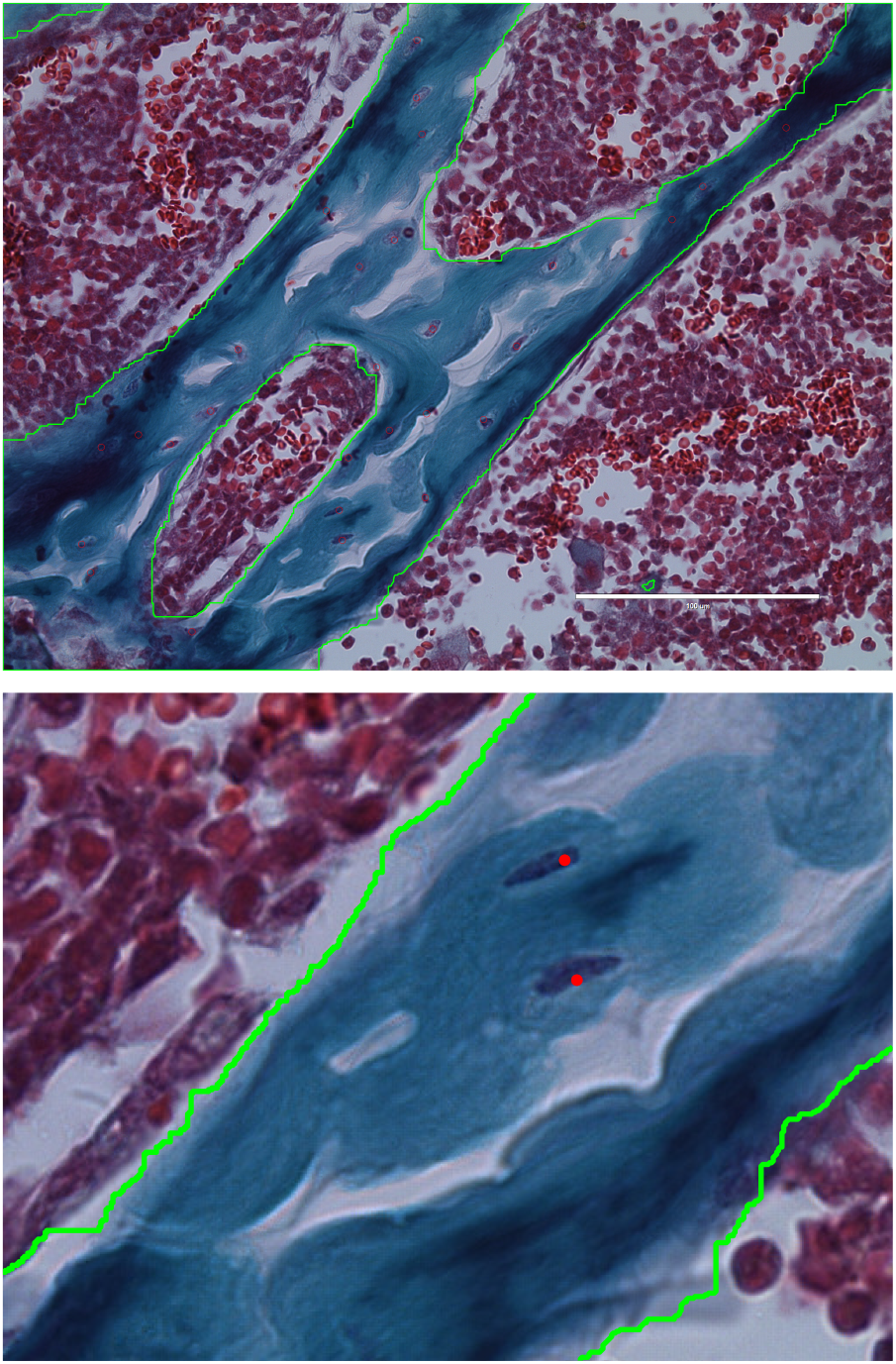
Output of image processing technique for osteocyte density calculation: (top) full size image, (bottom) close-up image. Region measured is outlined in green and osteocyte centers are marked in red.

Our criterion for what was marked and counted as an osteocyte was as follows:

i. The cell boundary of the osteocyte must be visible.
ii. There must be de-calcified osteoid around the osteocyte.

## B Monte Carlo Algorithm for Stochastic Simulation of Bone Formation

We give pseudocode for a fixed time step Monte Carlo algorithm for our osteocyte network formation model described in Section 1.2.1 using functions and parameters as detailed in Table 2 and mathematical details in Table 4. The Monte Carlo algorithm is broken town into two sub-algorithms. Algorithm 1 is the master algorithm that calls on Algorithms 2 and 3.

**Table 4.**
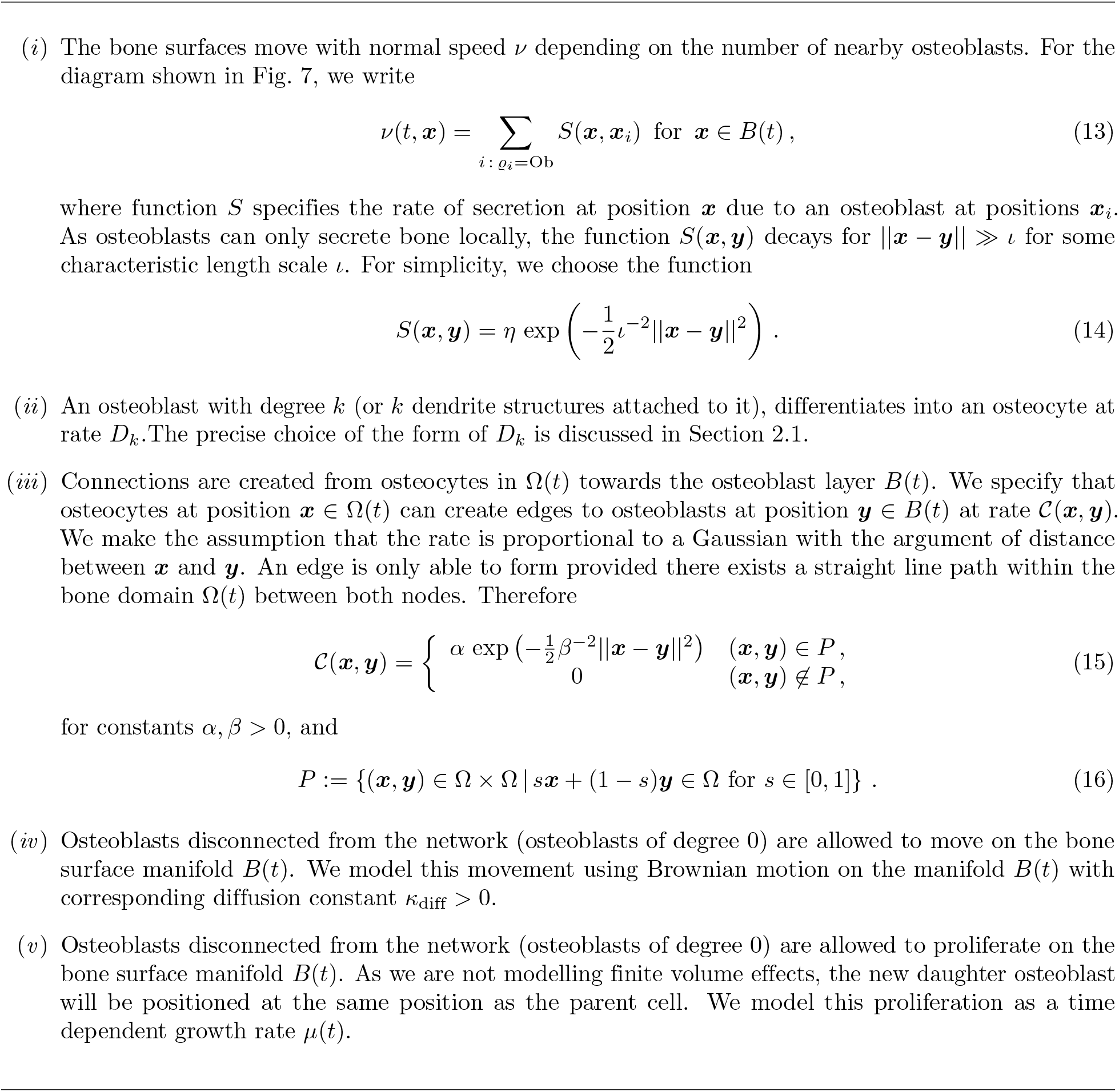
Model description: (*i*) bone secretion; (*ii*) osteoblast differentiation; (*iii*) dendrite growth; (*iv*) osteoblast migration; and (*v*) osteoblast proliferation.

Algorithm 2 details for how the network structure changes as the following Poisson processes: edge creation events from osteocyte to osteoblast; osteoblast-to-osteocyte differentiation; and osteoblast proliferation. Algorithm 2 follows from the Algorithm given in the appendix of [29].

Algorithm 3 details the changing domain Ω(*t*) and boundary *B*(*t*) (we specify osteoblast migration as part of this). For the changing domain, one has a few options regarding how one deals with the boundary (e.g., the level set method), but due to the fact that we have osteoblasts that occupy positions on the boundary, we use a particle method (discretising the boundary with an ordered set of particles).

As osteoblasts can diffuse on the boundary *B*(*t*), osteoblast diffusion is incorporated into this Algorithm 3 via a position jump process (a type of random walk) where osteoblasts can jump either left or right along the discretised boundary as a Poisson process that approximates diffusion (Brownian motion) as the spacing between the boundary discretisation reduces to zero. Therefore, osteoblasts move perpendicular to *B*(*t*), and secrete bone normal to Ω(*t*).

We now give extra details for Algorithm 3 on the particle method for the changing boundary and the selection of the jumping rate for the osteoblasts.

### B.1 Extra details for Algorithm 3

Algorithm 3 deals with the changing boundary by allowing for the addition and removal of extra discretisation points. When pairs of discretisation points are too close, one of the points may be removed to avoid having to reduce the time-step Δ*t*. When pairs of discretisation points move to far apart, a new discretisation point is added with coordinates equal to the mean of the pair of discretisation points in question.

There is the potential for topology changes where osteoblasts would be buried without differentiation. However, we do not account for topology changes as circular regions with an osteocyte within them would shrink to a point; we save on computation time by shrinking closed to loops to points immediately (see Figure 6).

**Fig 6.**
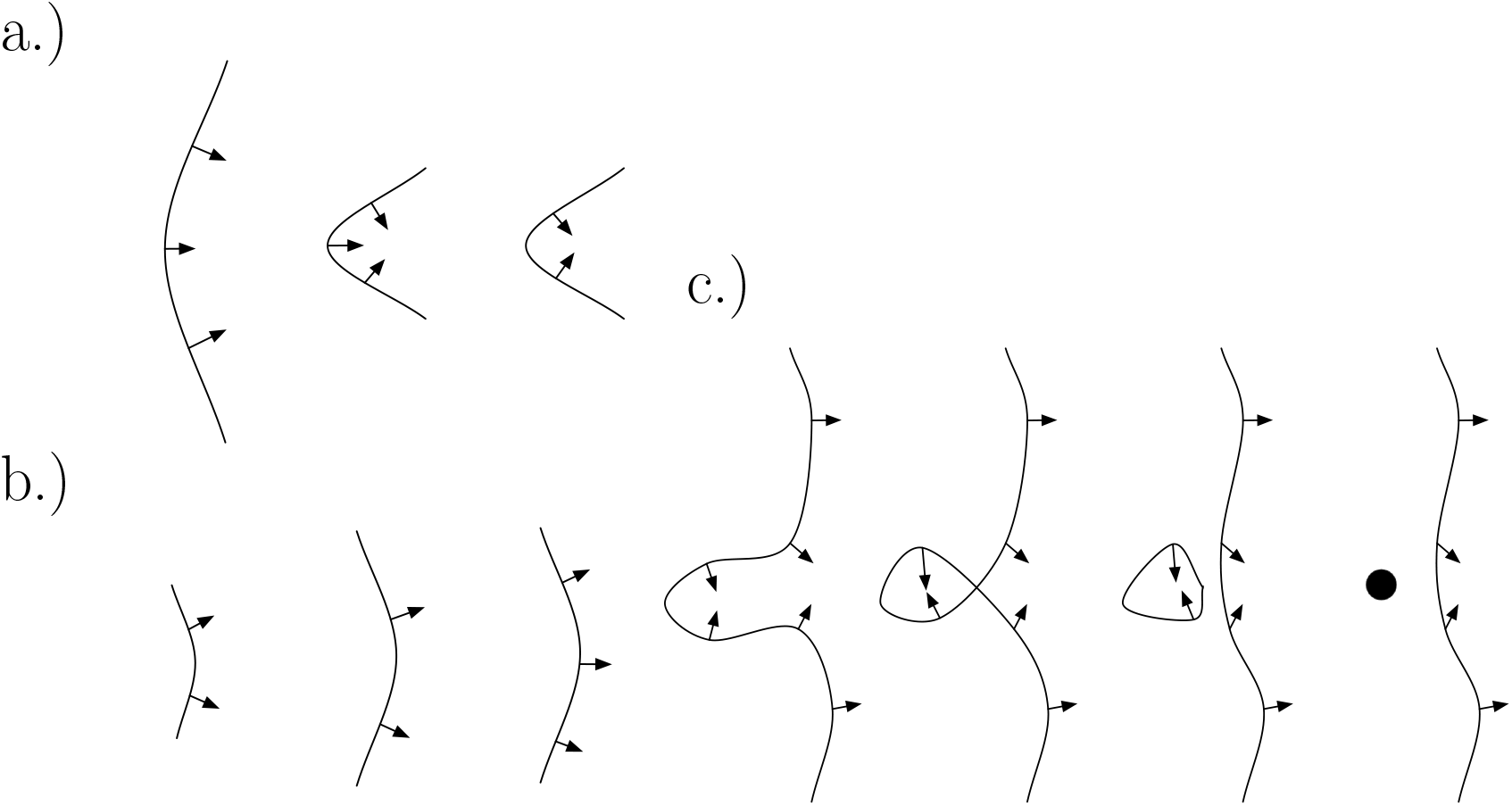
Diagrams showing boundary discretisation refinements: (a) shows a discretisation point being removed as the surrounding points move closer together; (b) shows a discretisation point being added as the surrounding points move further apart; and (c) shows how topology changes may occur, in the far right graphic, a hole is immediately collapsed (which saves on computation time).

Algorithm 3 approximates osteoblast diffusion via a jumping process along the boundary discretisation. The effective jump rates are found from analysing the Taylor expansion for the Poisson process where one either jumps “up” a distance of *ℓ_u_* at rate *j_u_* or “down” a distance of *ℓ_d_* at rate *j_d_* along an arc length coordinate *s*. As an approximation, we assume that the migration speed of osteoblasts occurs at an order of magnitude faster than the rate of osteoid deposition; therefore we can assume the interface is temporarily static. Consider a small time step of size Δ*t* > 0, and denote the density of osteoblasts on *B* by *ρ*(*t, s*), then

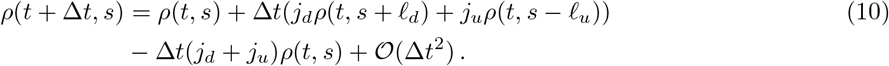

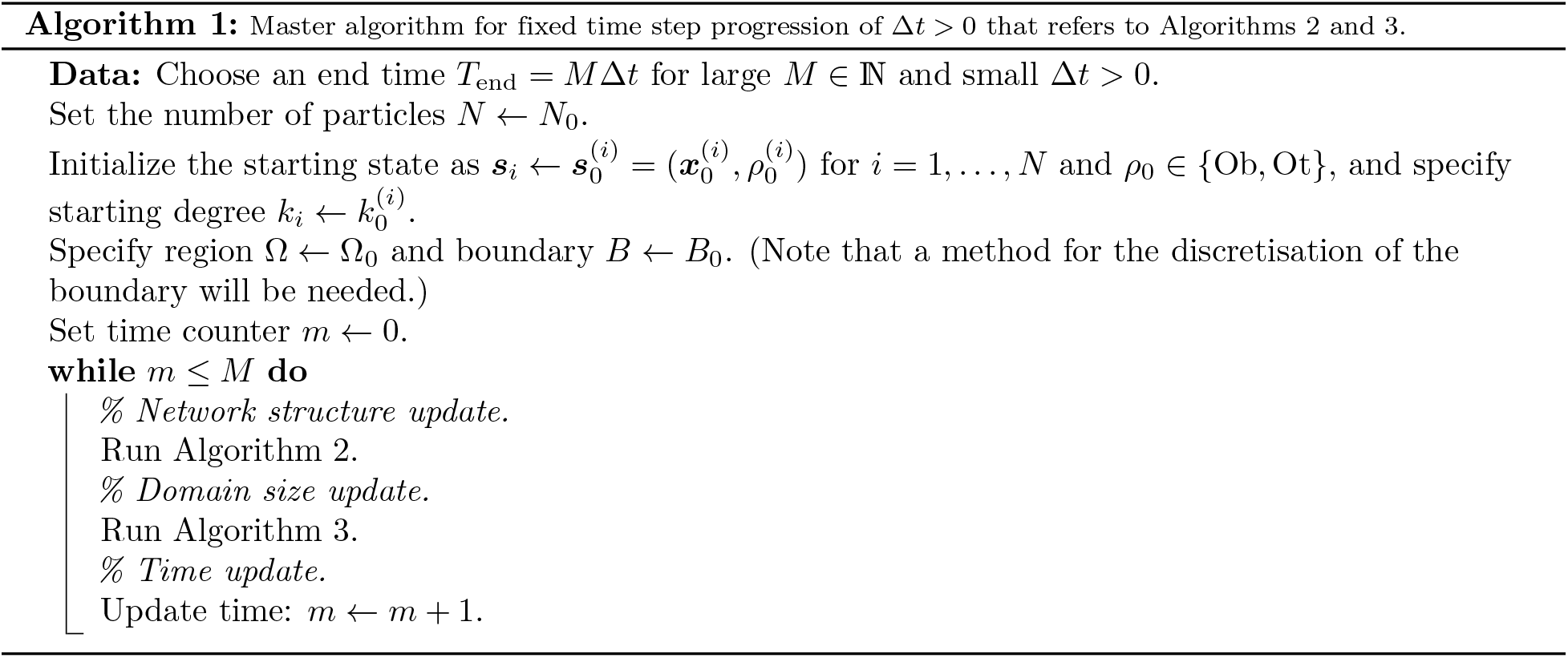

Taylor expanding around the value *s*, and taking the limit as Δ*t* → 0, we obtain the drift-diffusion equation

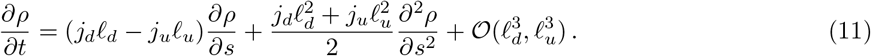

To prevent artificial drift of osteoblasts along the bone surface, one must choose *j_d_ℓ_d_* = *j_u_ℓ_u_* and we specify that the diffusion constant is 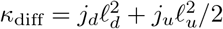. Therefore we select the jumping rates *j_u_* and *j_d_* as

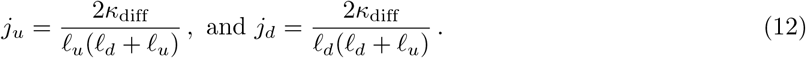

## C Mathematical description

We make the model description more precise through the introduction of mathematical concepts and notation. In our model there are two species: osteocytes and osteoblasts. These species act as nodes in a spatial network. Between pairs of nodes, undirected unweighted edges represent dendrites growing away from osteocytes. The osteocytes occupy an expanding domain 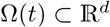, and osteoblasts are located on sections of the boundary of the domain *B*(*t*) ⊂ *∂*Ω(*t*). The boundary *B*(*t*) moves in the outward normal direction ***n***(*t*, ***x***) for ***x*** ∈ *B*(*t*) at velocity ***u***(*t*, ***x***). In Fig. 7, we give a schematic of a general domain.

**Fig 7.**
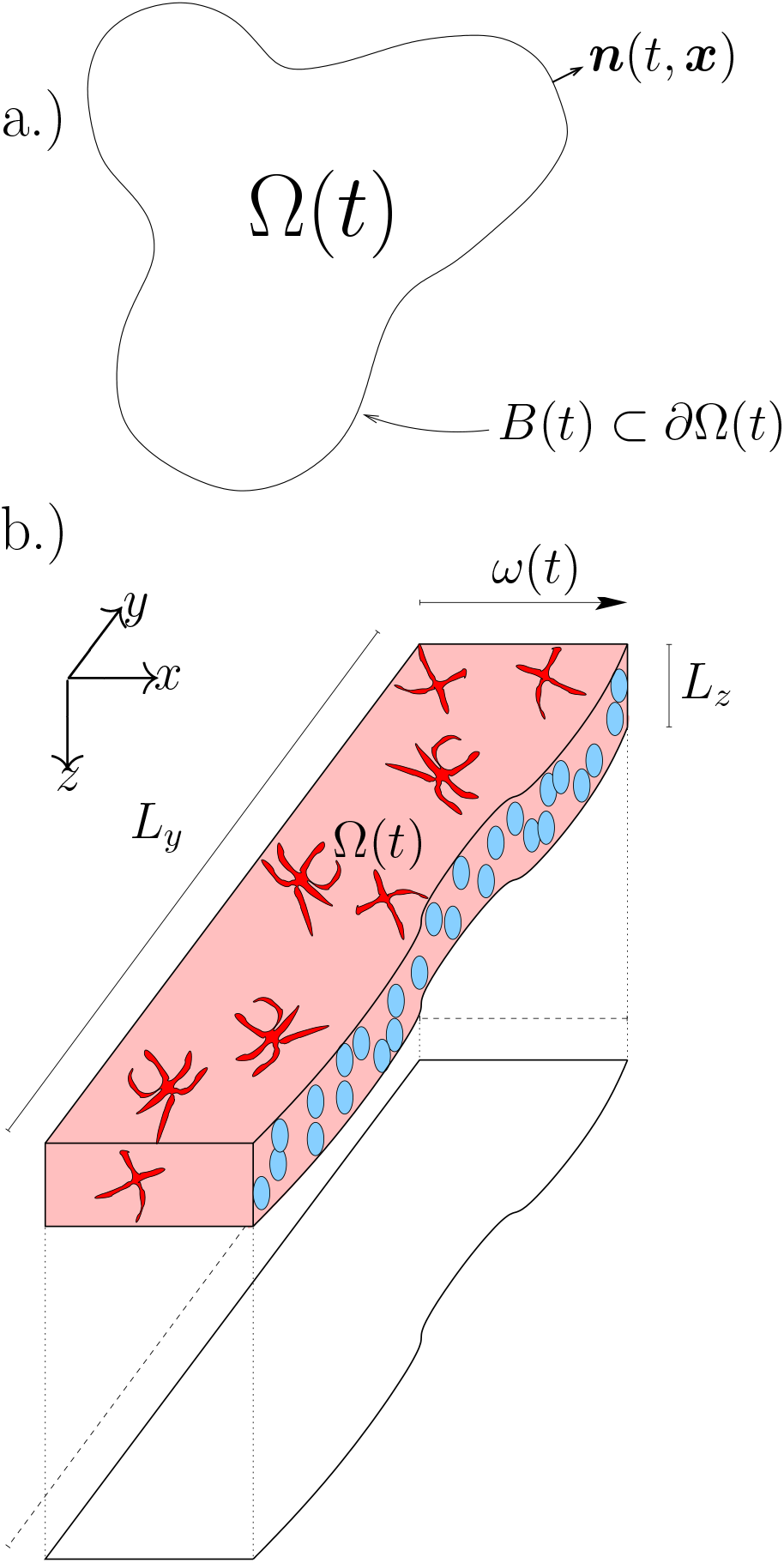
Domain diagrams for mode (a) general diagram of the domain Ω(*t*), and (b) domain specific to model realisations. Osteoblasts are depicted in light blue, osteocytes are shown in red. Two dimensional projection shown below.

For all times *t* > 0, the whole system is described completely by 4 (multidimensional) quantities. First, one must know how many nodes *n* are present. Second, we specify the cell type *ϱ_i_* for each node *i* = 1,…, *n*. We must know whether it is an osteoblast (Ob) or an osteocyte (Ot). Third, each node *i* = 1,…, *n* must have an associated position, we write 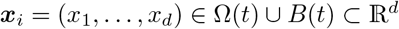. Finally, we must know the adjacency matrix of the underlying network. The adjacency matrix of a network is a matrix 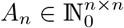 with entries that detail the number of unweighted undirected edges between nodes *i* and *j*. Our framework [29] allows for the existence of multiedges (multiple edges between the same two nodes) for mathematical tractability.

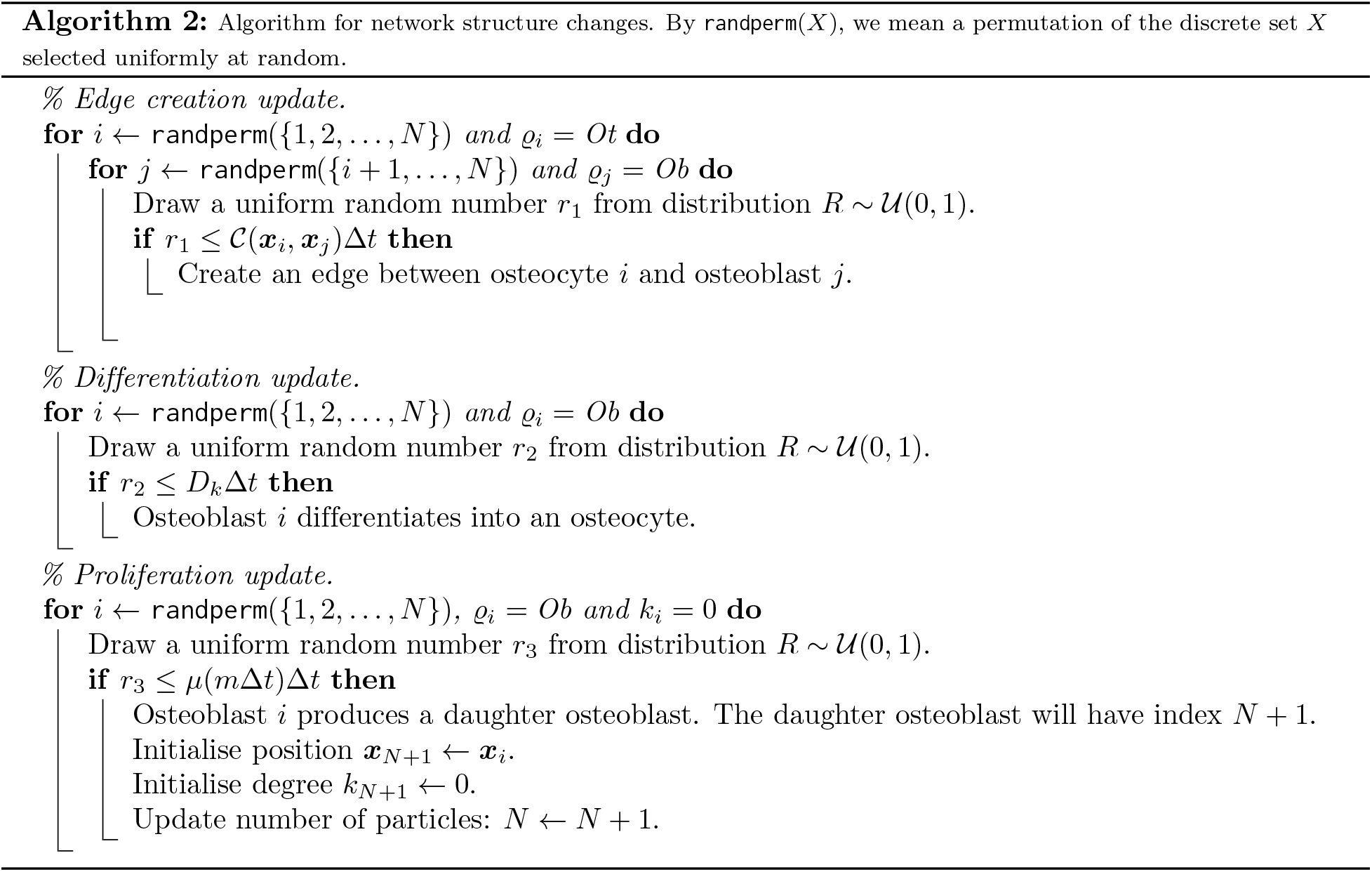

Although as the size of the network becomes large, the probability of multiedges occurring approaches zero. Additionally, the mean degree for each node is *a priori* likely to be low, e.g., we used 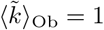.

We now respecify the model given in Figure 2 (and mathematical in Table 4). We frequently make use of the phrasing that *events occur at some rate*. This rate will always then refer to the probability of an event happening per unit time (also known as a Poisson process).

We list the functional forms underpinning our model in Table ??; in the case of the rate of osteoblast differentiation (*D_k_*), we investigate possible choices that represent different dependences on network structure. Recurring parameters that have fixed values used are listed in Table 2. Both Tables ?? and 2 contain brief details on how function forms/parameters were chosen; for extra details, see Appendix E.

### C.1 Derivation of mean-field equations

From earlier work [29], we can write down differential equations for the Local State Degree Distributions (LSDD). In Ref. [29], the expression *u_k_*(*t, **s***) was the expected number of particles with degree *k* and state vector ***s***. For the model contained in Section C, our state vector is described by position and cell type, so ***s*** = (***x***, *ϱ*), and *ϱ* ∈ {Ob, Ot}.

For notational convenience, we write *v_k_*(*t, **x***) is the expected number of osteoblasts of degree *k* with position ***x*** at time *t*, and *w_k_*(*t, **x***) as the expected number of osteocytes of degree *k* with position ***x*** at time *t*. We write the number density of osteoblasts and osteocytes (respectively) as

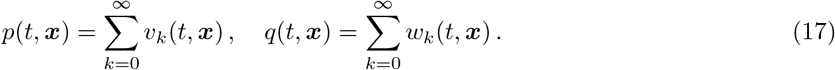

The osteocytes are present in domain 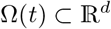, and the osteoblasts are present on (a subset of) the boundary of this domain *B*(*t*) ⊂ *∂*Ω(*t*). The domain boundary moves with normal velocity

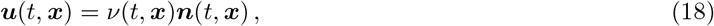

where ***n*** is the normal to *∂*Ω. Using Stone’s derivation of surfactants on moving interfaces [54], our equations become

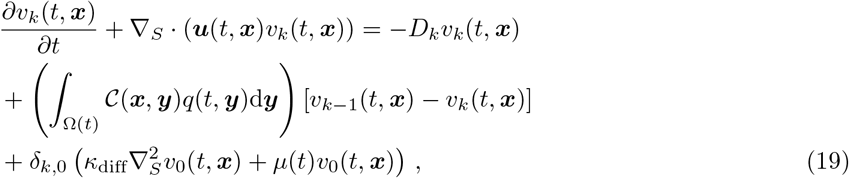

in *B*(*t*), and

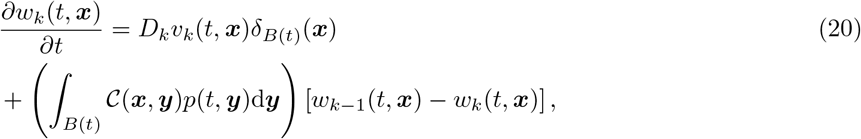

in Ω(*t*), where ∇_*S*_: = (*I_d_* − ***nn***^*T*^)∇ is the surface gradient operator^2^; Taking the continuum limit in equation (13), then

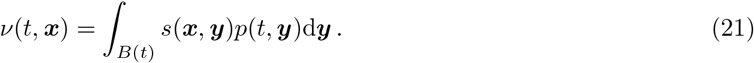

Assuming a solution homogenous in the *y* and *z* directions, we can write the system’s domain as: Ω(*t*) = (−∞, *ω*(*t*)]; the normal vector as ***n*** = ***e***_*x*_; and then

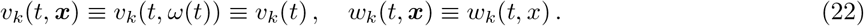

Under these assumptions, equations (19) and (20) become

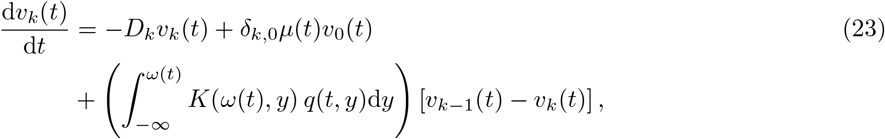

on the bone surface *B*(*t*) = {*ω*(*t*)} for kernel 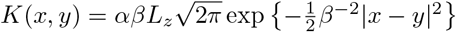, and

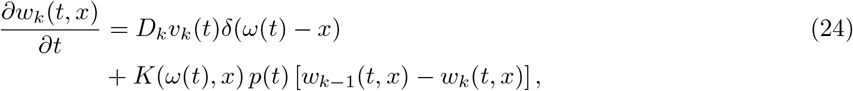

in the bone volume Ω(*t*). The velocity given by equation (21) is then interpreted as an ODE for the position of *ω*(*t*), which is

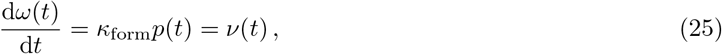

for secretion rate

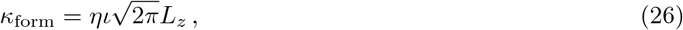

which represents the volume of bone secreted per osteoblast per unit time.

Through the use of the method of characteristics, we can write equation (24) as an integral of the solution of equation (23), see Appendix D.1 for full details. One can then use a standard finite differences numerical scheme to solve equation (23).

## D Solution method

We now need to solve equations (23), (24), and (25). We consider two different scenarios. In the first case (Section D.1), we turn the differential equations for *w_k_*(*t, x*) into integral equations involving the solution to *v_k_*(*t*) so that we only need to construct a numerical scheme for the ODEs for *v_k_*(*t*). In the second case (Section D.2), we choose the rate of proliferation *μ* to conserve mass [See equation (48)], allowing a travelling wave solution which can be solved analytically.

### D.1 Time dependent problem

Analysing equations with a moving boundary can be challenging, so we change our coordinate system to a fix the location of this boundary by moving from (*x, t*) to (*z, τ*), where *z* = *x* − *ω*(*t*) and *τ* = *t*. Therefore we write equation (24) as

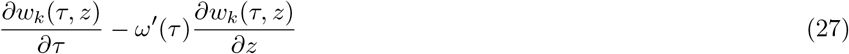

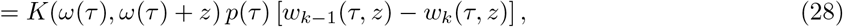

on 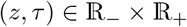 with initial condition

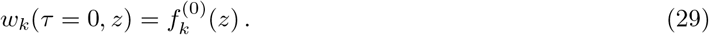

Additionally, we turn the delta function in equation (24) into the boundary condition

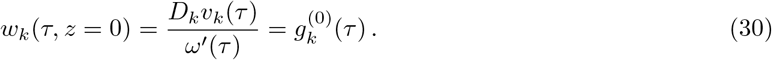

We use the Method of Characteristics using *ζ* as the characteristic variable, we find

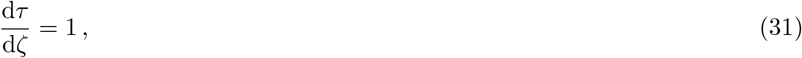

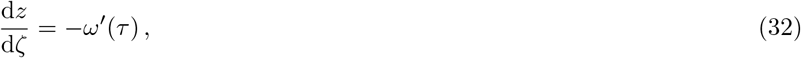

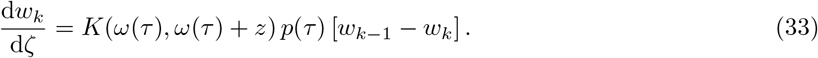

When *ζ* = 0, we specify initial and boundary data as a function of *ξ*. We write

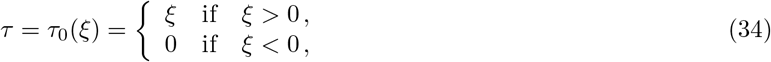

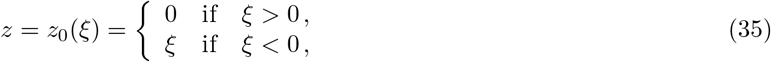

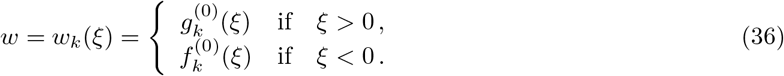

Solving equations (31) and (32), we find that

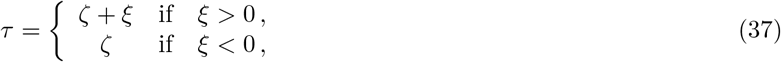

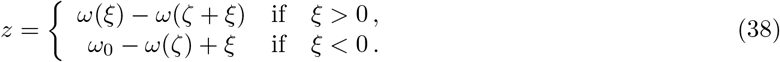

Writing

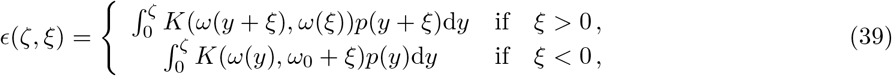

then

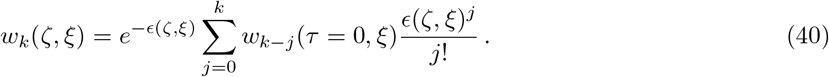

Inverting equations (37) and (38), we obtain

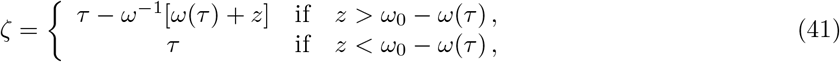

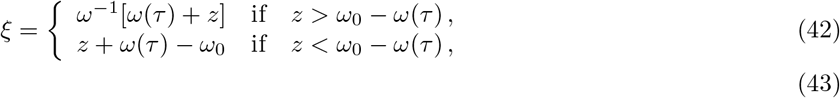

and therefore the solution as a function of (*z, τ*) is given by inserting equations (41) and (42) into equation (40). For the exponent term *ϵ*(*z, τ*), we can use a change of variables and write

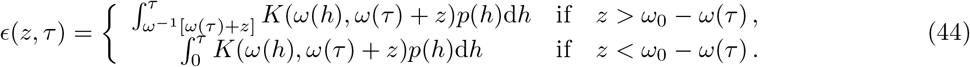

and we have also therefore solved our equations for *w_k_*(*t, x*). To solve equation (23) for *v_k_*(*t*), we need to evaluate the integral

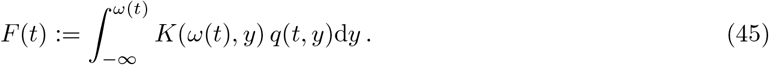

By noting that

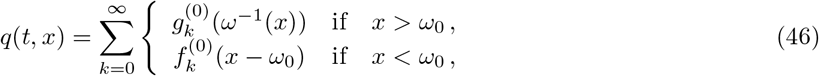

then we can calculate *F* as the contribution from the initial condition 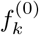 and the boundary condition 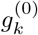, therefore

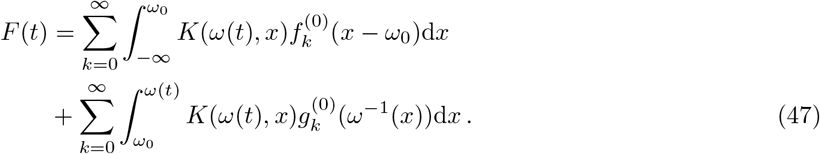

Referring back to the definition of 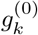 in equation (30), we note that our ODEs have become integro-differential equations. These can be solved via finite differences.

### D.2 Time independent problem: mass conserving travelling waves

To maintain a constant surface density of osteoblasts, we must choose

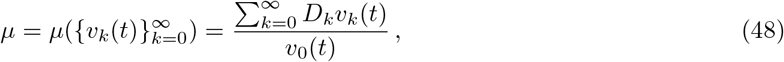

so that

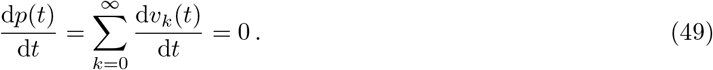

Our system of ODEs then admits a travelling wave solution corresponding uniform bone growth. We write the travelling wave speed as 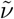, the solutions of the ODEs now correspond to the constant value 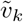 with sum 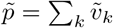. The solutions for the degree *k* osteocytes have travelling wave profile 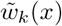 where *x* → −∞ corresponds to the back of the wave (away from the osteoblast front) and *x* = *ω*_0_ is the front of the wave (where the osteoblasts are located). The total osteocyte density 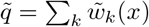 is then a constant by equation (46) as the initial condition 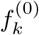 does not enter into the solution.

From equation (25), the wave speed is given by the relation

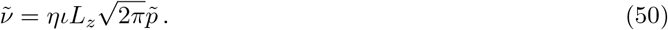

We now have to solve the algebraic system of equations

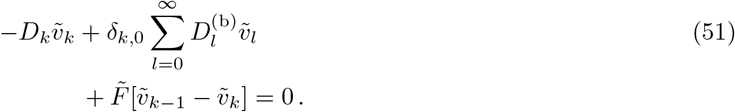

The constant 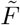 from equation (45) from is given by

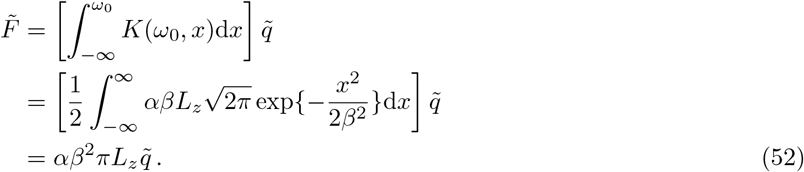

Summing equation (30) over *k* and using equation (50), 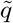 is given by

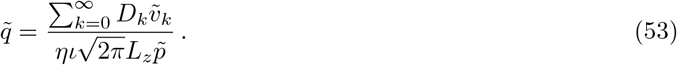

The mean osteoblast degree can be found by solving equations (51) and calculating 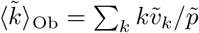. We can also calculate the solution to 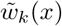 by using equation (40) and assuming steady-state bone formation, that is

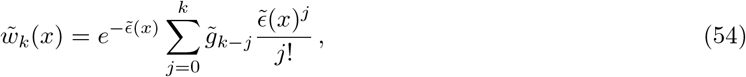

where

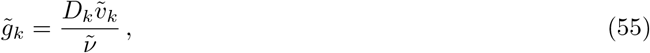

and

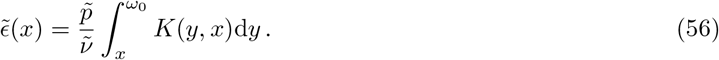

With a solution to 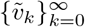, then the mean osteocyte degree of connectivity can be calculated away from the osteoblast front as

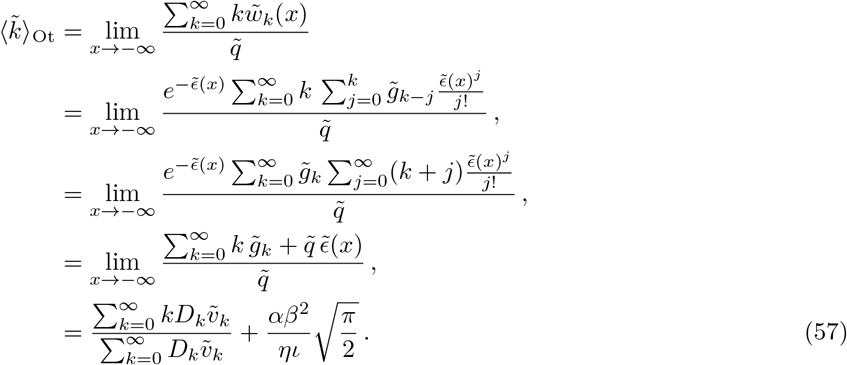

We then use equation (57) for different choices in *D_k_* to aid with parameterisation.

#### D.2.1 No network influence: Null hypothesis

Rewriting equation (5) below

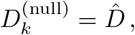

one can insert this choice of *D_k_* into equations (51)–(53) to show that the osteocyte number density is

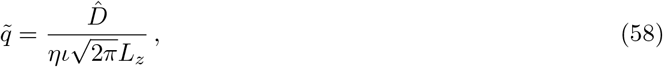

the mean osteoblast degree is

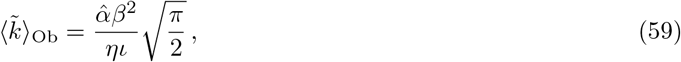

and the mean osteocyte degree of connectivity is

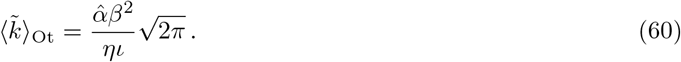

Quite remarkably, it is always the case that 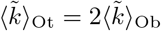.

#### D.2.2 Proposed mechanism: Switch-like influence

Rewriting equation (8),

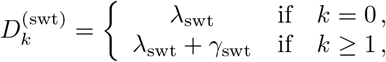

one can insert this choice of *D_k_* into equations (51)–(53) to show that

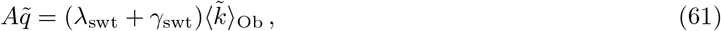

for *A* = *α*_swt_*β*^2^*πL_z_* and

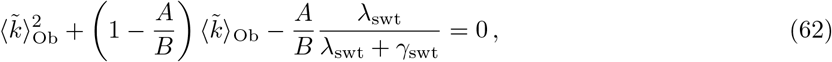

for 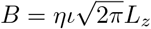. The solutions of the equations above are algebraically tumultuous — so we do not present these here! Additionally, the mean osteocyte degree of connectivity is

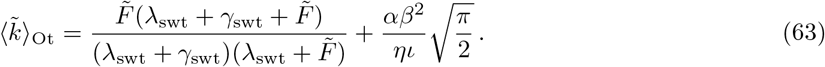

Note there is consistency that the solutions to equations (61)–(63) agree with the solutions to equations (58)–(60) when *γ*_swt_ = 0.

## E Parameter estimation

### E.1 Osteoblast diffusion constant (*κ*_diff_)

In the work by Araujo *et. al*. [23], a cellular automata model was proposed in which migrating osteoblasts were allowed to move on a lattice every time step. With time steps of 6 mins, in the absence of any chemical signalling, osteoblasts moved either up, down, left or right with equal probability. Additionally, it was stated that (from experiments), the speed of a migrating osteoblast was 0.1470 mm day^−1^.

Using the results from Taylor-King *et. al*, it was reported for a velocity jump process in *n*-dimensions with fixed speed *S_T_*, exponentially distributed runs of mean length *μ_τ_*, and with uniformly random selection of new directions, the large time effective diffusion constant is

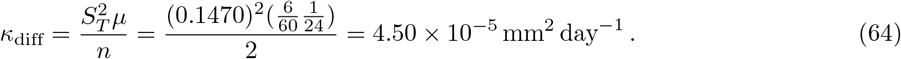

### E.2 Domain size (*L_z_, L_y_*)

*L_y_* is chosen such that *L_y_* ≫ *β. L_z_* is chosen to be the diameter of an osteoblast (15 × 10^−3^ mm) plus the mean distance between osteocytes (25 × 10^−3^ mm). Also note that *ω*_0_ = 0

### E.3 Bone secretion rate and shape parameters (*η, ι*)

Through analysis of the travelling wave case (see Appendix D.2), one can choose *η* such that one obtains the (experimentally observed [23]) travelling wave speed of 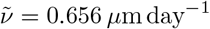. This is done by noting that for a constant wave speed 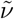, by equation (25), we choose that

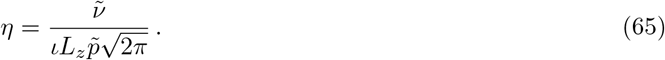

The shape parameter *ι* is essentially the standard deviation of the normally distributed bone secretion function and is chosen to be the typical diameter of an osteoblast.

### E.4 Dendrite growth shape parameter (*β*)

This value is essentially the standard deviation of the normally distributed rate function [Equation (15)]. Note that we are not selecting the length of dendrites with this parameter, it specifies a how likely a dendrite is likely to form from an osteocyte within the bone to an osteoblast on the bone surface. The length of the dendrite may change while the osteoblast is located on the bone surface, once it differentiates into an osteocyte, the edge length is fixed.

We choose *β* to be the typical distance between osteocytes. The typical distance measured between osteocytes was reported in Ref. [30] to be between 10–40 *μ*m; we use the midpoint in this range.

### E.5 Osteoblast and osteocyte densities 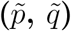

Osteoblast surface densities are observed within the range 2–10 × 10^3^ mm^−2^, the value of p was chosen to be the midpoint in this range. Osteocyte densities are observed within the range 1.90–2.85 × 10^4^ mm^-3^, the value of 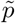 was chosen to be the midpoint in this range. Both of these values are used in steady-state travelling wave regime.

### E.6 Mean osteoblast degree 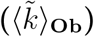

The only network structure property that we deduced from experimental data was the average number of distinct osteocytes that a single osteoblast connects to. This corresponds to the average node degree of osteoblasts in our connectivity network, 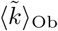. We assumed this number to be 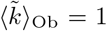 based on the following estimate. We divide the total number of dendritic processes of an osteoblast, by the redundancy of the physiological network, i.e. by the number of dendritic processes connecting the same osteoblast and osteocyte. Kamioka *et. al*. [16] measured the number of dendritic processes coming from an osteoblast to be 4.8. The redundancy of osteocyte–osteoblast connections can be estimated by dividing the number of processes running from an osteocyte to osteoblasts (26.05) by the number of osteoblasts reached by a single osteocyte (5.7), giving a redundancy of 4.57 [16]. This provides the estimate 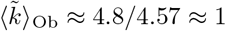. This value is consistent with Kamioka *et. al*.’s observation that they rarely observed osteoblasts that had connections with more than one osteocyte. We note here that there are not many studies reporting these kinds of measurements. Marotti *et. al*. [51] have reported the number of dendrites coming from an osteoblast to range from 9.4 to 20.9 in four osteoblasts (so ~ 13.6 on average). If this figure is used instead, 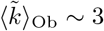.

## F Discussion on *D_k_*

### F.1 Proposed mechanism: Cumulative activation

One mechanism we considered was a cumulative activation effect, essentially each osteoblast has an network-independent rate of differentiation (λ_1_) plus an network-dependent contribution (*f* = *γ*_1_*k*). This network-dependent contribution corresponds to increasing the rate of osteoblast-to-osteocyte differentiation proportional to the number osteocytes that each osteoblast is in contact with. We write

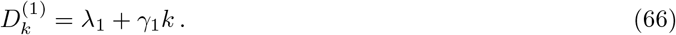

One can calculate the mean osteoblast degree with the following analysis. Writing the first moment of 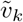 as 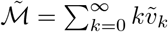, then the equation for *k* = 0 is

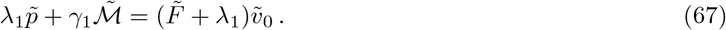

We can recursively find all 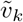, and we determine that

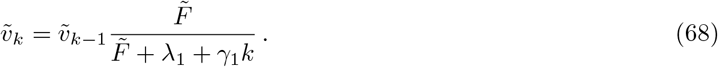

though the use of lower incomplete gamma functions defined as 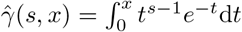, one can show that

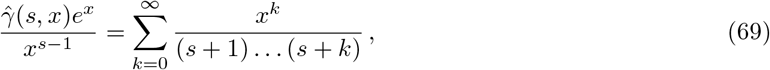

and therefore

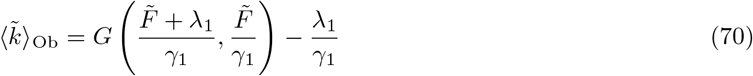

for 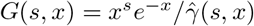 and

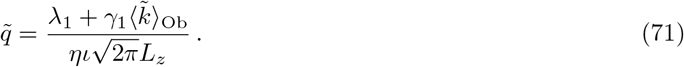

We fit parameters as stated in Section 2.1.2. Specifying that the network has an excitatory effect on osteoblast differentiation, we set 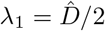 and therefore *α*_1_ = 2.41 × 10^−3^ day^−1^ and *γ*_1_ = 1.30 × 10^−3^ day^−1^. Figure 8 shows that when considering the cumulative activation model when compared to the null model, it takes approximately 3 years to get to the desired osteocyte density of 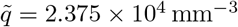. This timescale is clearly too long to be considered biologically realistic. Additionally, were one to make *γ*_1_ < 0, the rate 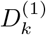 can be negative which means that when using this model, inhibitory network effects are impossible. When considering the cumulative activation model, at the time of osteoblast differentiation a large quantity of a differentiation-promoting protein must be present in the osteoblasts. This protein would have therefore previously diffused along the dendrite structures. From a biological perspective, it seems more likely that only a small amount of protein should be required to travel though the dendrite structure to induce differentiation.

**Fig 8.**
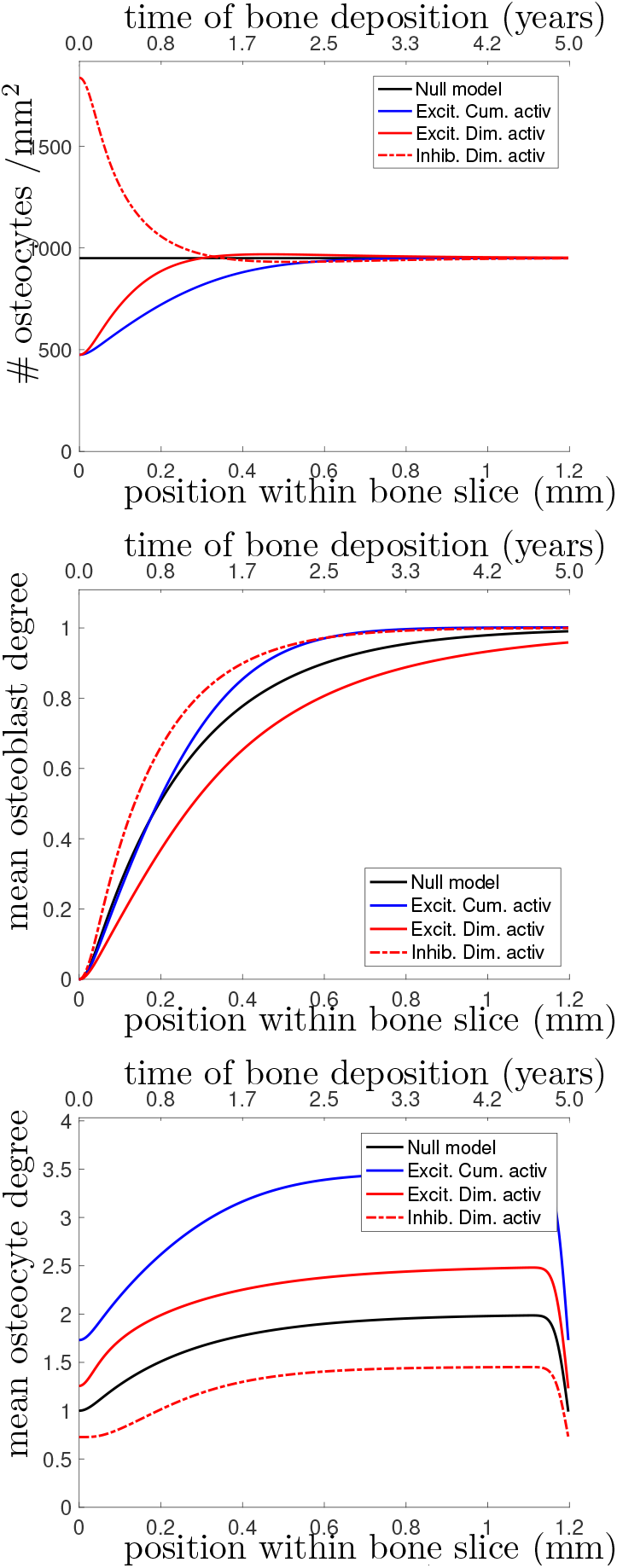
(Top) Osteocyte density profile 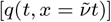, (Middle) mean osteoblast degree over time 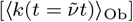, and (Bottom) mean osteocyte degree over time 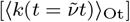 when solving equations (19)–(20). The black line shows the null model, the blue line shows the excitatory cumulative activation model, and the red lines shows the diminishing activation model, excitatory effects using the solid line and inhibitory effects using the dot-dashed line.

### F.2 Proposed mechanism: Diminishing activation

Another mechanism considered is a cumulative inhibition effect. Each osteoblast has an network-independent rate of differentiation (λ_2_) plus an network-dependent contribution (*f* = *γ*_2_/*k*). This network-dependent contribution corresponds to decreasing the rate of osteoblast-to-osteocyte differentiation inversely proportional to the number osteocytes that each osteoblast is contact with. We write

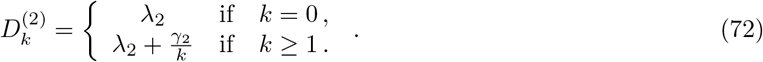

To calculate the mean osteoblast degree, one can make use of the the series

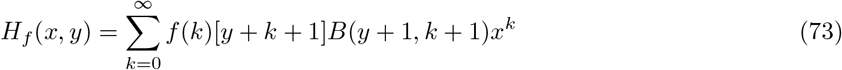

for 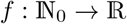 and 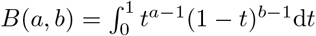 is the beta function to calculate

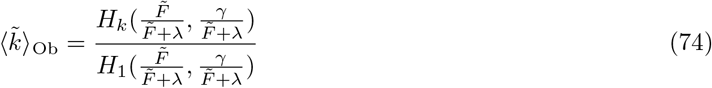

and

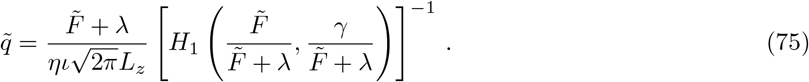

Using equation (52), algebraic equations (74)–(75) are closed.

We fit parameters as stated in Section 2.1.2. Specifying that the network has an excitatory effect on osteoblast differentiation, we set 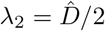 and therefore *α*_2_ = 1.75 × 10^−3^ day^−1^ and *γ*_2_ = 4.56 × 10^−3^ day^−1^. Our diminishing activation model also has the possibility to allow for inhibition when 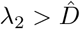: however the rate 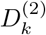 can be negative. For the greatest allowable amount of inhibition, one can set *γ*_2_ = −λ_2_. To maintain the desired travelling wave profile, the maximum value of λ_2_ is given as 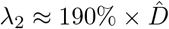 and in which case *α*_2_ = 1.01 × 10^−3^ day^−1^.

Both the excitatory and (maximum) inhibitory effects are shown in Figure 8. When compared to the null model, both models take approximately 2 years to get to the desired osteocyte density. This timescale is too long to be considered biologically realistic.

One emergent property we notice from this choice in differentiation mechanism is that when *D_k_* is non-monotonic (i.e., has a maximum/minimum around *k* = 1), it appears that there is a over-reaching then adjustment period when viewing the density profile *q*(*t, x*), e.g., the density starts below 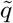, then over-reaches above 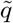 before converging to 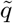. When viewing 3 dimensional scans, one does observe similar behaviour around cement lines [55, 56].

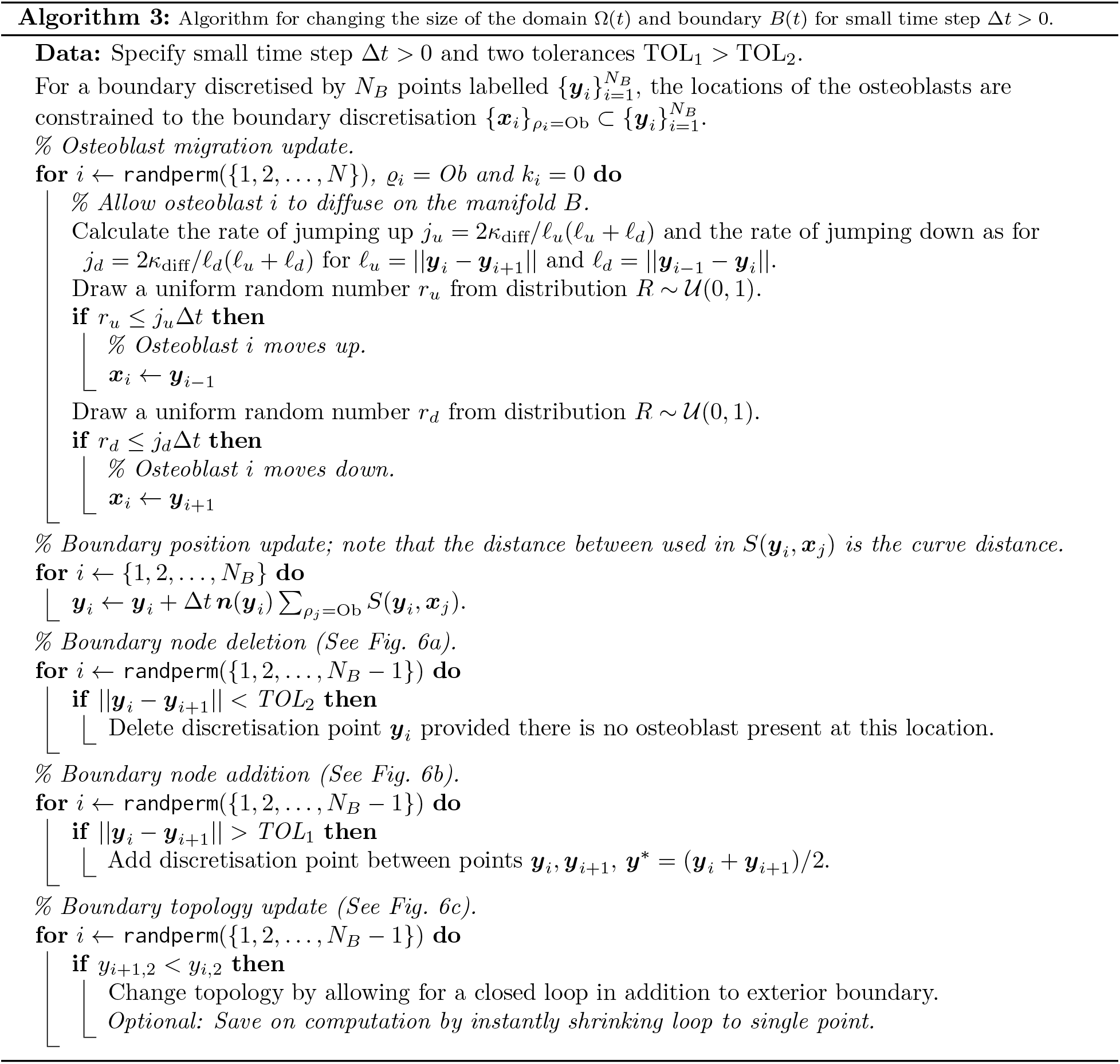

**Table 5.**
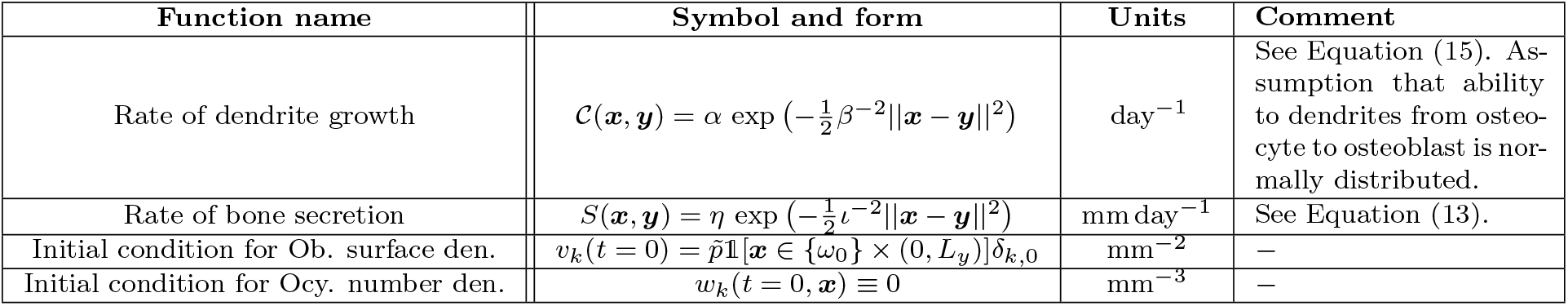
Model functions.

**Table 6.** Choice in rate of osteoblast differentiation.

| Symbol and form | Interpretation |
| --- | --- |
| $D_k^{(\text{null})} = \hat{D}$<br>This corresponds to the null hypothesis: the network does not impact the rate of osteoblast differentiation. | No network effects. |
| $D_k^{(\text{swt})} = \begin{cases} \lambda_{\text{swt}} & \text{if } k = 0, \\ \lambda_{\text{swt}} + \gamma_{\text{swt}} & \text{if } k \geq 1. \end{cases}$<br>The network does impact osteoblast differentiation rates but there are no cumulative effects, i.e., communicating with more osteocyte does not mean osteoblasts differentiate any faster/slower. | Switch-like influence. |
| $D_k^{(1)} = \lambda_1 + \gamma_1 k$<br>The osteoblast will differentiate at a rate that increases proportional to how many osteocytes it is in communication with. | Cumulative activation. |
| $D_k^{(2)} = \begin{cases} \lambda_2 & \text{if } k = 0, \\ \lambda_2 + \gamma_2/k & \text{if } k \geq 1. \end{cases}$<br>The osteoblast will differentiate at a rate that increases inversely proportional to how many osteocytes it is in communication with. | Cumulative inhibition. |

## Footnotes

1 These include Receptor Activator of Nuclear Factor Kappa-B Ligand (RANKL), Vascular Endothelial Growth Factor (VEGF) [11], Parathyroid Hormone (PTH) [13], calcium ions (Ca^2+^) [14], and Sclerostin [15].

2 We use standard convention that *I_d_* is the *d*-dimensional identity matrix.

